# A Role for Importin α in Ciliogenesis and Cilia Length Regulation during Nephrogenesis

**DOI:** 10.1101/2025.04.10.648263

**Authors:** Natalie Mosqueda, Patrick James Sutton, Christopher W. Brownlee

## Abstract

Cilia are evolutionary conserved microtubule-based organelles involved in many biological processes. Motile cilia facilitate fluid flow, while primary cilia transduce developmental and growth signals. Defects in cilia structure or function can lead to a spectrum of ciliopathies. Ciliogenesis and length regulation depend on protein transport to the ciliary base, but the molecular mechanisms remain unclear. We demonstrate that the nuclear transport protein, importin α, when membrane-localized via palmitoylation, or non-palmitoylated in the cytoplasm, localizes to the base or lumen of primary and motile cilia. There we find it is required to initiate ciliogenesis and maintain proper cilia length. We found importin α binds Cep164, Cep78, and Arl13b, proteins essential for ciliogenesis and length maintenance. Disrupting importin α palmitoylation in *Xenopus laevis* lead to abnormal kidney morphology and reduced renal primary cilia. These findings reveal a novel pathway through which importin α contributes to ciliogenesis and advance understanding of ciliopathy etiology.

## Introduction

Cilia are specialized organelles that protrude from the plasma membrane and are composed of a microtubule-based axoneme encased by a ciliary membrane. Most cells possess either a single immotile primary cilium or one or more motile cilia. Primary cilia play critical roles in normal development and tissue homeostasis by transducing signals from the extracellular environment. These signals encompass morphogens, growth factors, light perception, and mechanosensation^1^. Multiciliated epithelial cells contain hundreds of motile cilia, which are essential for generating fluid flow within the brain ventricles, spinal cord, and reproductive and respiratory tracts^2^. Defects in cilia structure or function can lead to a class of human genetic disorders known as ciliopathies. There are currently 35 known ciliopathies and 187 ciliopathy associated genes that affect multiple organs in the body^3,4^. Many ciliopathies are associated with defects in the structural components required for ciliogenesis, the complex process which describes the assembly of cilia. One important component of ciliogenesis is the basal body, which is comprised of the mature mother centriole. The mother centriole contains distal and subdistal appendages that are essential for docking to the plasma membrane. Once the basal body docks at the plasma membrane, a cholesterol-rich membrane termed the ciliary pocket gives rise to the formation of the transition zone^1,5^. The transition zone acts as a selective filter for regulating trafficking into and out of the cilium along the axoneme in order to maintain proper length and signal transduction functions. Despite numerous studies investigating these processes, the precise molecular mechanisms underlying ciliogenesis and cilia length maintenance remain unknown.

One interesting pattern that has emerged from various studies investigating ciliogenesis and cilia length maintenance is the discovery that several proteins involved in nuclear transport localize within the cilia and are essential for cilia formation^6^. The karyopherin transport protein family, known as importins, comprise the primary nuclear import proteins. This family includes importin α, with seven human isoforms (KPNA1-7) and importin β, which has 2 isoforms (KPNB1-2)^7,8^. While importin β_2_ can bind cargo directly through conserved PY motifs, importin β_1_ requires heterodimerization with importin α to bind cargo. Importin α functions as an adapter protein, binding cargo in the cytoplasm via nuclear localization signal (NLS) sequences^8,9^. The cargo bound to importins can subsequently traverse the nuclear pore by passing through the nuclear pore protein (NUP) selective filter, after which the cargo is released upon the binding of importin β to RanGTP. Notably, importin β_2_, RanGTP and three NUPs have all been identified as localizing to the primary cilia and are required for ciliogenesis^10–13^.

Recent work has unveiled novel functions for importin α when localized to the plasma membrane via post-translational lipid modification known as palmitoylation^14,15^. This phenomenon was initially identified by Brownlee and Heald 2019, which demonstrated that importin α isoform 1 (KPNA2), when modified with four palmitate lipids, is partitioned to the cell cortex where it serves as a key regulator of nuclear and spindle size^15^. More recently, it has been demonstrated that the residues of importin α found to be palmitoylated in *Xenopus laevis* (*X. laevis*) are also conserved in humans. This same study further revealed a role for palmitoylated importin α in regulating mitotic spindle orientation^15^. Building upon this previous work, which highlight the involvement of palmitoylated importin α in orchestrating membrane-bound processes, along with the presence and necessity of other nuclear transport proteins during cilia formation, we propose a hypothesis in which palmitoylated importin α may mediate ciliogenesis and cilia length maintenance through the binding of NLS-containing ciliary cargo proteins.

The current work demonstrates through confocal and ultra-expansion microscopy (U-ExM) the characterization of importin α localization in cilia and at the cilium base in both in vitro human primary cilia and in vivo *X. laevis* epidermal multiciliated cell (MCC) models. We show that the palmitoylation status of importin α and its ability to bind and release cargo proteins is essential for both primary and motile ciliogenesis. Furthermore, we highlight the critical requirement for a balance between palmitoylated membrane-bound and non-palmitoylated cytoplasmic importin α for cilia length maintenance in both primary and motile cilia. We also offer a bioinformatic analysis identifying proteins known to be involved in cilia biology which we predict to contain a NLS sequence and using this list identify the ciliary proteins Cep78, Arl13b, and Cep164 as binding partners of importin α that are required for proper regulation of cilia length and cilia formation. Lastly, we explore how disruption of importin α palmitoylation leads to decreased primary cilia in the kidney as well as a kidney ciliopathy phenotype in *X. laevis* embryos. Overall, this work offers a previously unexplored role for palmitoylated importin α in ciliogenesis and length maintenance in both motile and primary cilia and advances our understanding of the etiology of ciliopathies.

## Results

### Importin α localizes to ciliary structures in human epithelial primary cilia and *X. laevis* epidermal MCCs

Importin β_2_ has previously been identified as having a role in trafficking proteins to primary cilia and NUPs have been shown to localize to the cilia base^10–12,16^. This suggests that importin mediated transport may be playing a role in ciliary functions. Consequently, we aimed to investigate the role of importin α in ciliogenesis by first characterizing its localization to ciliary structures. To achieve this, hTERT-RPE-1 (RPE-1) cells were probed for importin α ciliary localization, as this cell line is a well-established model system of ciliogenesis^5,17^. RPE-1 cells were ciliated after two days of serum starvation and imaged by deconvolution epifluorescent microscopy which revealed localization of importin α throughout the primary cilia, with significant enrichment at the cilium base (**Figure 1A**). To further elucidate its localization at the base, RPE-1 cells were labeled with a basal body associated protein, Chibby, which we found to colocalize with importin α at the cilium base (**Figure 1B**). Given the diminutive size of basal bodies (centriolar width and length are approximately 250nm and 400nm, respectively^18–20^), we employed U-ExM for increased resolution of ciliary importin α localization. U-ExM is a technique that enables the enlargement of biological samples for enhanced visualization and resolution of preserved ultrastructures^21,22^. To demonstrate the feasibility of this technique, we first performed U-ExM on ciliated RPE-1 cells and successfully expanded the sample, allowing for the visualization of more intricate staining for the primary cilium and basal body (**Figure 1C**). Notably, U-ExM imaging clarified the localization of importin α throughout the basal bodies of RPE-1 cells, corroborating our previous observations of importin α at the cilium base (**Figure 1D**). To determine whether this localization is conserved within motile cilia, we utilized the *X. laevis* model system. The *Xenopus* system is an ideal model for studying in vivo ciliogenesis, as the developing *Xenopus* embryo contains MCCs along the entire outer body axis, known as epidermal MCCs^23,24^. The *X. laevis* epidermal MCCs are an advantageous model to study ciliogenesis due to their rapid development and external localization on the epidermis, facilitating easy manipulation and visualization^24,25^. Immunostaining of epidermal MCCs from a *X. laevis* NF stage 30 embryo revealed importin α localization throughout motile cilia and at the cilia base, consistent with our findings in RPE-1 cell primary cilia (**Figure 1E**). Another unique aspect of the *Xenopus* system is the ability to physically shear and collect motile cilia from the epidermal MCCs. This process, termed reversible deciliation, involves a calcium-shock to the embryo, causing the cilia to shed while the basal bodies remain at the apical surface followed by subsequent cilia regrowth^25^ (**Figure 1F**). By employing this technique, we aimed to biochemically characterize the presence of importin α in isolated cilia. Motile cilia from epidermal MCCs were isolated from *X. laevis* NF stage 30 embryos and immunoblot analysis revealed the presence of importin α in the isolated motile cilia validating our microscopy results (**Figure 1G**). Collectively, these findings demonstrate through a multifaceted approach, the conserved localization of importin α at ciliary structures.

**Figure 1:**
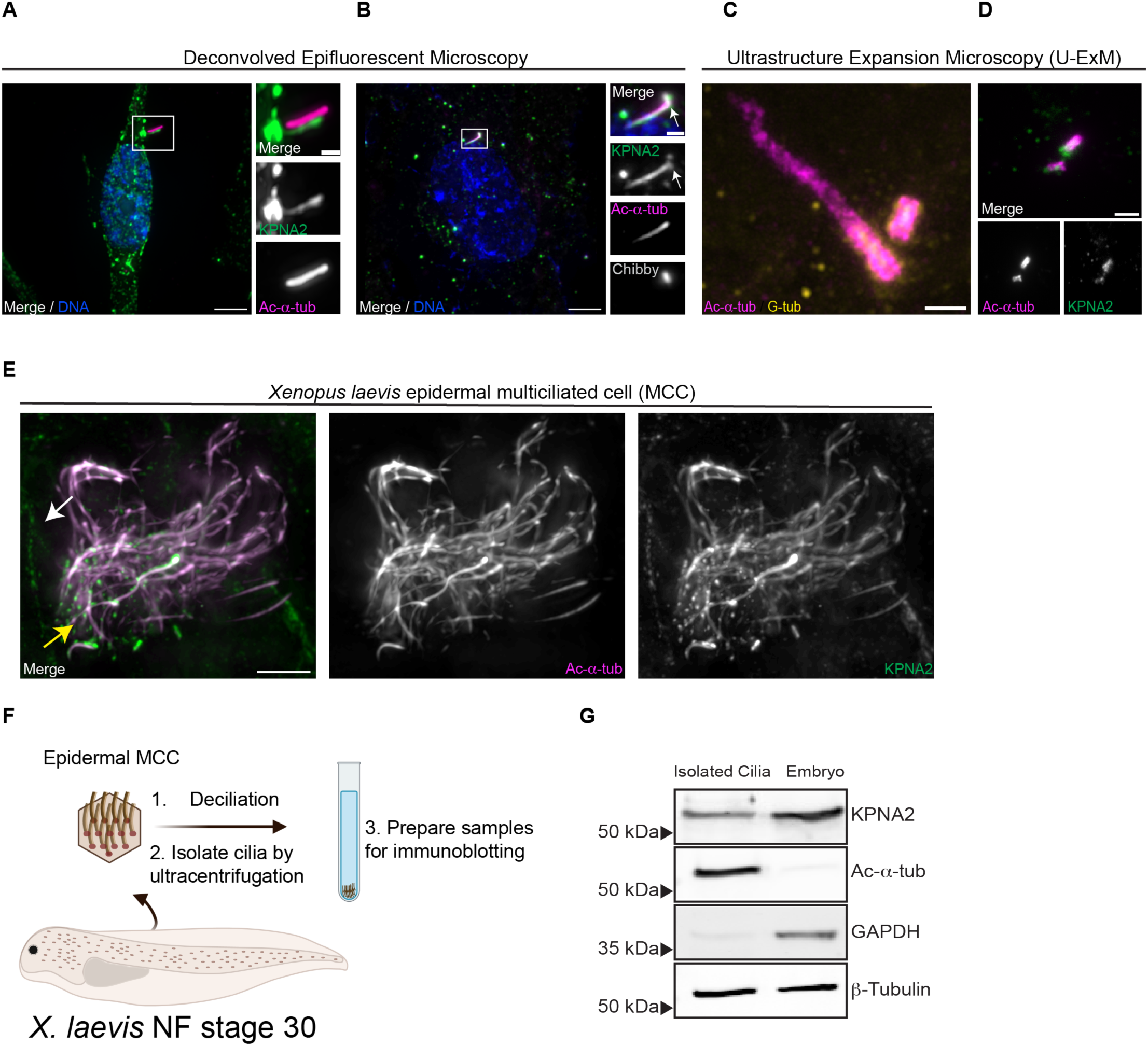
Importin α localizes to primary cilia in human epithelial RPE-1 cells and Xenopus laevis (X. laevis) epidermal multiciliated cells (MCCs). To induce ciliogenesis, RPE-1 cells were serum starved for 48 hr and immunostained to identify importin α localization. **A)** Deconvolved epifluorescent image of an RPE-1 cell immunostained for KPNA2 (importin α) (green), acetylated-α-tubulin (magenta), and DNA (blue). Scale bars = 5µm (full image) and 1µm (inset image). **B)** Deconvolved epifluorescent image of an RPE-1 cell immunostained for KPNA2 (importin α) (green), acetylated-α-tubulin (magenta), Chibby (gray) and DNA (blue). Scale bars = 5µm (full image) and 1µm (inset image). **C)** Ultra-Expansion confocal microscopy (U-ExM) image of a ciliated RPE-1 cell immunostained for acetylated-α-tubulin (magenta) and gamma-tubulin (yellow). Scale bar =2 µm. **D)** U-ExM images of ciliated RPE-1 cells immunostained for KPNA2 (importin α) (green) localization with acetylated-α-tubulin (magenta). Scale bar = 2µm. **E)** Deconvolved epifluorescent image of X. laevis NF stage 30 epidermal MCCs immunostained to characterize KPNA2 (importin α) (green) localization at cilia, acetylated-α-tubulin (magenta), the plasma membrane (white arrow) and at the cilia base (yellow arrow). Scale bar = 5µm. **F)** Schematic of deciliation protocol on X. laevis NF stage 30 embryos. **G)** Immunoblot of isolated cilia and whole NF stage 30 X. laevis embryo for KPNA2 (importin α), acetylated-α-tubulin, GAPDH, and β-tubulin.

### Palmitoylated importin α regulates ciliogenesis in RPE-1 cells

Previous research has demonstrated that importin α can be targeted to the plasma membrane upon palmitoylation, thereby regulating processes such as nuclear and spindle size scaling, and more recently mitotic spindle orientation^14,15^. Given our characterization of importin α at ciliary structures, we hypothesized that membrane localization of importin α via palmitoylation, may play a regulatory role in ciliogenesis. To investigate this, we initially explored the potential role of palmitoylated importin α in regulating ciliogenesis in RPE-1 cells, which form primary cilia via the intracellular ciliogenesis pathway resulting with its hallmark ciliary pocket (**Figure 2A**). The intracellular pathway begins with the mother centriole acquiring Golgi apparatus-derived vesicles containing proteins which begin the assembly of the ciliary sheath internally, before migrating apically to the plasma membrane, creating the ciliary pocket that serves as a platform for vesicular trafficking^26,27^. Once the basal body docks at the plasma membrane, the cilium will begin to protrude outwardly. To ascertain whether membrane association of importin α via palmitoylation regulates ciliogenesis, pharmacological inhibitors were used to disrupt the addition and removal of palmitate lipids, palmitoylation and depalmitoylation respectively. S-palmitoylation and O-palmitoylation are the processes of attaching palmitate lipids to cysteine or serine residues, respectively, through either the action of palmitoyl acyl transferase (PAT) or membrane-bound O-acyltransferases (MBOAT) enzymes. In this study, we used the inhibitor Wnt-C59, a specific small molecule inhibitor of porcupine O-acyltransferase (PORCN), which is responsible for serine palmitoylation of only two known targets: Wnt, whose palmitoylation is required for its secretion^28^, and importin α, whose palmitoylation is essential for its membrane localization^14^. The effect of Wnt-C59 results in the decrease of palmitoylated membrane-bound importin α and an increase in unpalmitoylated cytosolic importin α (**Figure S1A**). Depalmitoylation, however, involves the removal of palmitate lipids, regulated by the enzyme class acyl protein thioesterases (APTs). To target depalmitoylation we used the inhibitor palmostatin which inhibits the activity of APT 1/2, broadly inhibiting the depalmitoylation of palmitoylated proteins, leading to an increase in palmitoylated membrane-bound importin α and a subsequent decrease in cytosolic importin α (**Figure S1A**). Additionally, importazole was used to inhibit the ability of importin α to release NLS-containing cargo^29^.

**Figure 2:**
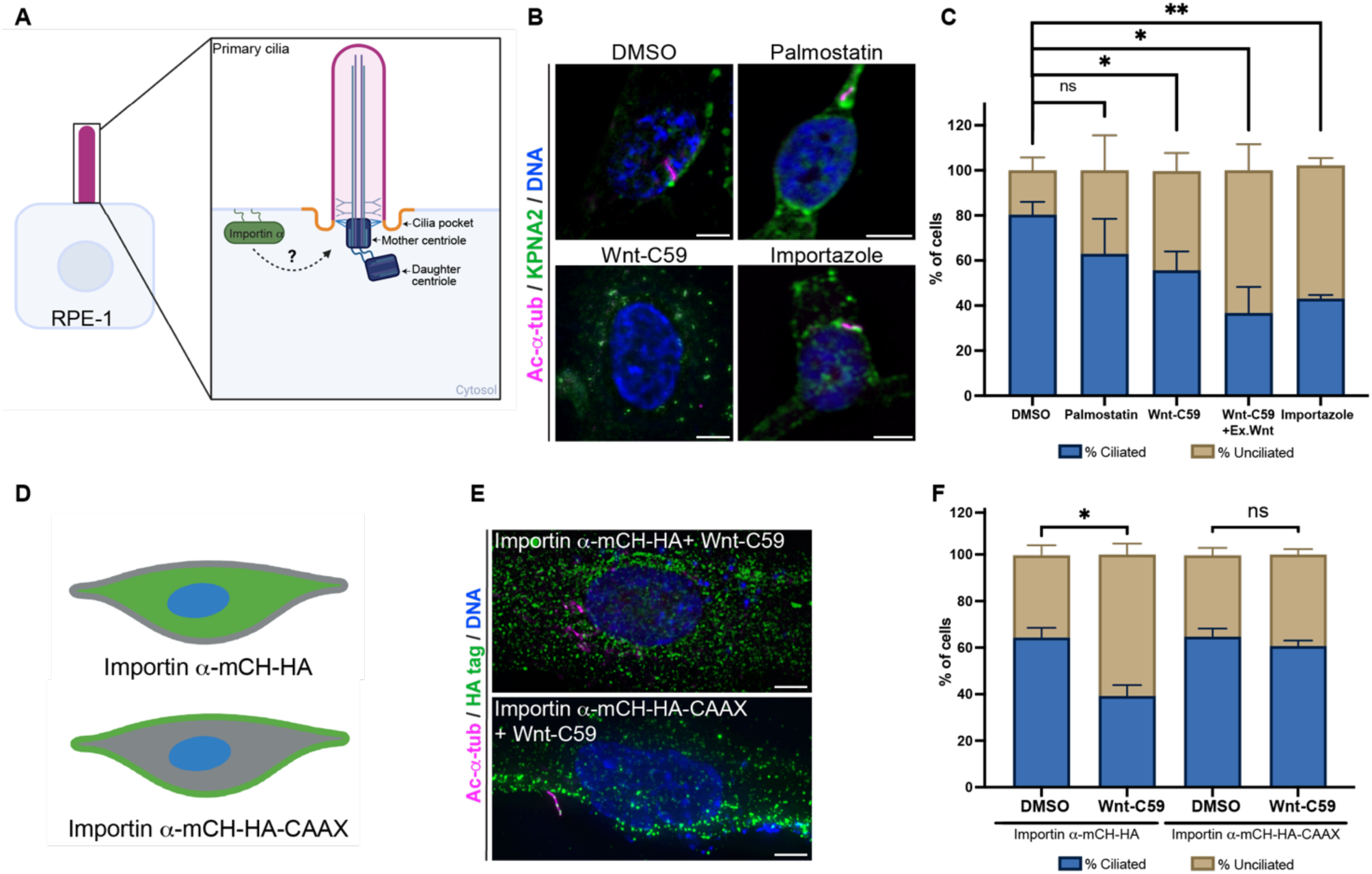
Palmitoylated importin α regulates primary ciliogenesis of human epithelial RPE-1 cells. **A)** Schematic of hypothesized role for palmitoylated importin α regulating the formation of primary cilia in RPE-1 cells. RPE-1 cells form cilia via the intracellular pathway as shown with its hallmark ciliary pocket on both sides of the cilium (orange). **B)** Non-ciliated RPE-1 cells were serum starved for 24 hours and treated with listed the drugs for an additional 24 hours in serum free conditions: DMSO, 10μM Wnt-C59, 50μM palmostatin, or 40μM importazole. Deconvolved epifluorescent images of drug treated RPE-1 cells immunostained for KPNA2 (importin α) (green), acetylated-α-tubulin (magenta), and DNA (blue). Scale bar = 5µm. **C)** Quantification of %of ciliated vs. % of unciliated RPE-1 cells after drug array treatment for 24hrs.* P<0.05, ** P<0.01, Student’s t-test, n=300 cells per treatment, 3 replicates. Mean ± SEM. **D)** Schematic of RPE-1 cells transfected via nucleofection, with 1µg of either importin α-mCH-HA or importin α-mCH-HA-CAAX plasmid. **E)** Deconvolved epifluorescent images of RPE-1 cells transfected with importin α constructs, serum starved and treated with 10μM Wnt-C59 for 24 hours. **F)** Quantification of % of ciliated vs. % of unciliated RPE-1 cells transfected with either importin α-mCH-HA or importin α-mCH-HA-CAAX, serum starved and treated with DMSO or 10μM Wnt-C59 for 24hrs .* P<0.05, Student’s t-test, n=100 cells per treatment, 3 replicates. Mean ± SEM.

To elucidate the role of importin α in ciliogenesis, RPE-1 cells were serum starved for 48 hours to allow cells to enter G0 and induce ciliogenesis^1^. After 24 hours of serum starvation, RPE-1 cells were treated with the aforementioned drugs while remaining in serum free media for an additional 24 hours (**Figure S1B**)^1,30,31^. This experimental window was used to determine if palmitoylated importin α is required for cilia formation by quantification of the percentage of ciliated versus unciliated RPE-1 cells. Immunofluorescent analysis of RPE-1 cells treated with palmostatin revealed increased membrane-bound palmitoylated importin α, but no apparent effect on cilia formation (**Figures 2B and 2C**). Wnt-C59 treatment of RPE-1 cells resulted in a reduction of membrane-bound palmitoylated importin α, translocating it to the cytoplasm and concomitantly decreasing the percentage of ciliated cells (**Figures 2B and 2C**). To address any Wnt signaling off-target effects of PORCN inhibition by Wnt-C59, RPE-1 cells were additionally treated with Wnt-C59 under exogenous Wnt-conditioned media, which did not rescue the percentage of ciliated cells. Additionally, siRNA-mediated knockdown of PORCN in RPE-1 cells phenocopied the decreased ciliation results observed with Wnt-C59 treatment (**Figures S1C-S1E**). Knockdown of PORCN in RPE-1 cells resulted in a ∼15% ciliation reduction compared to control siRNA, and, intriguingly, ∼40% of the ciliated cells exhibited a stubby cilia phenotype, suggesting a potential effect on regulation of proper cilia length. (**Figures S1D and S1E**). Lastly, RPE-1 cells treated with importazole also exhibited a decrease in ciliogenesis, suggesting that importin α releasing bound NLS-containing ciliary cargos is essential for ciliogenesis to occur (**Figures 2B and 2C**). To determine if the observed ciliogenesis defects after Wnt-C59 treatment are attributable to the absence of membrane-bound importin α, we transfected RPE-1 cells with an importin α-mCH-HA-CAAX plasmid, which facilitates membrane localization via farnesylation through the CAAX motif, irrespective of palmitoylation status (**Figure 2D**)^15,32,33^. As outlined in the experimental timeline in Supp. Figure 1A, 24 hours prior to ciliogenesis, RPE-1 cells were transfected with importin α-mCherry-HA-CAAX (importin α-mCH-HA-CAAX) or a wild-type importin α plasmid (importin α-mCH-HA), followed by drug treatment with DMSO or Wnt-C59 to elucidate the requirement of membrane-bound importin α for proper ciliogenesis. Strikingly, we observed a rescue of ciliogenesis with importin α-mCH-HA-CAAX transfection in comparison to importin α-mCH-HA + Wnt-C59 and Wnt-C59 treatment alone (**Figures 2C and 2E-2F**). Lastly, as RPE-1 cells form cilia specifically via the intracellular pathway we explored if there is a conserved role for palmitoylated importin α in cells which utilize a different ciliogenesis program termed the alternative pathway. The alternative ciliogenesis pathway involves the basal body docking directly to the plasma membrane, after which there is a recruitment of ciliary proteins involved in building the cilium^26,34,35^. We wanted to test if palmitoylated membrane-bound importin α may also regulate this ciliogenesis pathway by acting as a protein scaffold to bind NLS-containing cilia proteins required for cilia formation. To test the importance for palmitoylated importin α in regulating the alternative pathway, canine renal MDCK-II cells were utilized. These cells were treated prior to ciliogenesis with the same array of drugs and monitored for changes in ciliation. MDCK-II ciliation results corroborated those observed in Figure 2C, demonstrating the requirement for membrane localization of importin α and its proper function to release bound cargo in the alternative ciliogenesis pathway (**Figures S1F and S1G**). Taken together, our data demonstrates a novel role for palmitoylated importin α regulating ciliogenesis.

### Palmitoylated importin α regulates ciliogenesis in *X. laevis* epidermal MCCs

Multiciliogenesis is a process involving centriole duplication that generates hundreds of basal bodies, which subsequently dock at the membrane and facilitate vesicle transport to deliver proteins required for cilia formation^24^. To investigate whether palmitoylated importin α regulates the distinct ciliogenesis pathway for the formation of multiciliated cells in the *X. laevis* epithelium (**Figure 3A**), *X. laevis* embryos at NF stage 18, the developmental stage characterized by the amplification of basal bodies, were treated with palmostatin or Wnt-C59 to inhibit importin α depalmitoylation and palmitoylation, respectively. *X. laevis* epidermal MCCs are present at NF stage 28, however *X. laevis* embryos were drug treated until NF stage 37 to ensure maximum drug penetration into MCCs (**Figure 3B**). Only the inhibition of importin α palmitoylation by Wnt-C59 treatment resulted in a decrease of the amount of MCCs per unit area in *X. laevis* embryos, suggesting a global effect on ciliated MCCs (**Figure 3C**). We also observed in *X. laevis* embryos that were treated with Wnt-C59 a shortened body axis phenotype previously ascribed to defects in the Wnt signaling pathway involving the development of the anterior-posterior axis^36,37^ (**Figure 3B**). Previous studies have demonstrated that various mucociliary cells on the epidermis rely on the Wnt/β-catenin signaling pathway for proper cell differentiation and patterning of mucociliary epithelium, although the timing of this signaling during development is unclear^23^. Therefore, we sought to elucidate whether the reduction in the quantity of epidermal MCCs, as evidenced by the absence of cells containing acetylated α-tubulin, is attributable to a lack of cellular differentiation of MCCs or a defect in ciliogenesis. We also wanted to test whether the effects on ciliogenesis observed upon PORCN inhibition were due to defects in the Wnt pathway, or if these defects are due to disruption of importin α membrane localization. To accomplish both of these objectives, we microinjected *X. laevis* embryos at the specific blastomere identified as giving rise to the epidermis with an importin α-mCH-HA or importin α-mCH-HA-CAAX plasmid, followed by drug treatment to determine if importin α membrane localization can rescue ciliogenesis defects. *X. laevis* embryos microinjected with importin α-mCH-HA-CAAX exhibited a rescue in ciliogenesis, as quantified by the intergraded density of the MCCs labeled for cilia with acetylated-α -tubulin (**Figures 3D and 3E**). In support of this result, we also disrupted PORCN expression by microinjecting a morpholino targeting PORCN and observed loss of cilia formation in the epidermal MCCs further demonstrating the role for palmitoylated importin α in regulating ciliogenesis (**Figure S2**). These results highlight the critical and conserved role of palmitoylated importin α in cilia formation of both primary cilia and multicilia, across in-vivo and in-vitro models.

**Figure 3:**
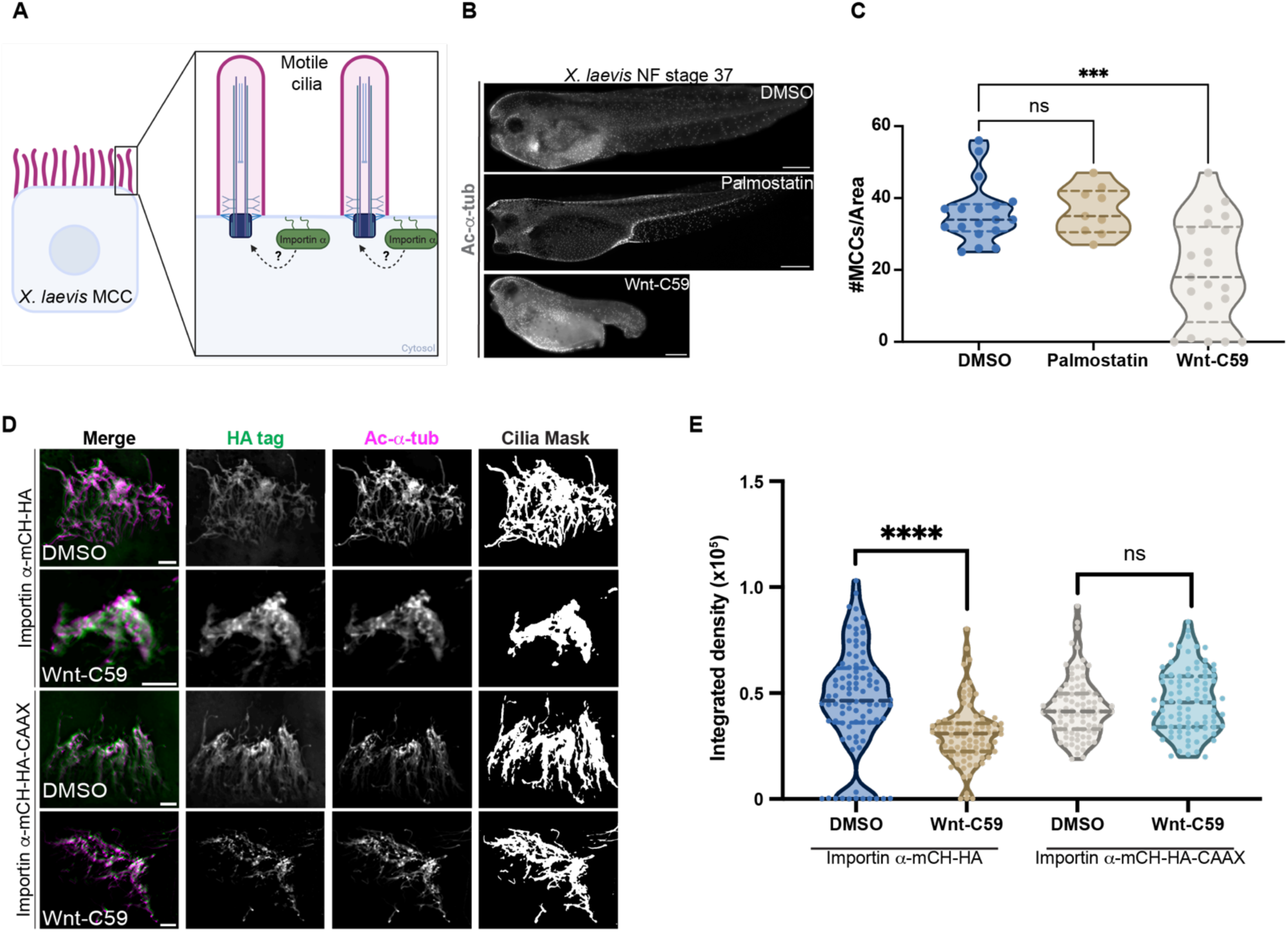
Palmitoylated importin α has a conserved role in regulating multiciliogenesis in *X. laevis* epidermal MCCs. **A)** Schematic of hypothesized role for palmitoylated importin α regulating the formation of motile cilia in the *X. laevis* epidermal MCC. **B)** *X. laevis* NF stage 37 embryos treated for 3 days with DMSO, 100µM Wnt-C59 or 1mM palmostatin followed by immunostaining of epidermal MCCs by acetylated-α-tubulin (gray). Scale bar = 200µm. **C)** Quantification of ciliated MCCs/area per drug treatment. ***P<0.001 Student’s t-test n=30 tadpoles. **D)** Deconvolved epifluorescent image of *X. laevis* embryos microinjected at the 4-cell stage with 30pg of importin α-mCH-HA or importin α-mCH-HA-CAAX targeted to epidermis. After microinjection with either plasmid, embryos were treated with either DMSO or 100 µM Wnt-C59 until NF stage 38. MCCs immunostained for acetylated-α-tubulin (magenta) and HA tag (green). Cilia masks were created by ImageJ mean thresholding of the acetylated-α-tubulin channel for quantification in Figure 3E. Scale bar = 5µm. **E)** Quantification of the integrated density in arbitrary units for each MCC cilia mask. **** P<0.0001 Student’s t-test, n=30 MCCs, 3 tadpoles, per condition.

### Importin α regulates cilia length in a palmitoylation and cytoplasmic dependent manner

Following ciliogenesis, the cilium maintains its structure through protein trafficking, ensuring the maintenance of its length and facilitating extracellular to intracellular signal transduction. Cilia length regulation is mediated by proteins trafficking into the cilium via the transition zone, enabling selective entry and exit of proteins in a bi-directional manner^38–41^. Previous studies have demonstrated the presence of a RanGTP gradient within cilia, in conjunction with the localization of nuclear pore proteins within these structures^10,11,13^. These findings imply that cilia may share fundamental characteristics with nuclear import, where protein trafficking is governed by the entry of importins through the selective nuclear pore protein complex and subsequent release upon interaction with RanGTP. Thus far our data has demonstrated that palmitoylated importin α regulates ciliogenesis and we hypothesize that importin α contains a dual role in ciliary maintenance by additionally regulating the steady-state dynamics of cilia length (**Figure 4A**). To assess for changes in cilia length, RPE-1 cells were serum-starved to induce ciliogenesis and then treated with the drug array to affect either palmitoylation status or importin function, once the cilia had already formed as measured by the CiliaQ ImageJ plug-in^42,43^ (**Figure S3A**). Previous work from our group has demonstrated that upon Wnt-C59 treatment, there is a shift of importin α membrane localization to the cytoplasm, while the converse is true upon palmostatin treatment (**Figure S1A**)^14,15^. Ciliated RPE-1 cells treated with Wnt-C59, inhibiting palmitoylation of importin α and thus increasing levels of cytoplasmic importin α, exhibited a decrease in cilia length (**Figures 4B and 4C**). Surprisingly, ciliated RPE-1 cells treated with palmostatin, which inhibits the removal of membrane localized importin α and therefore decreases the pool of cytoplasmic importin α, also resulted in shortened cilia (**Figures 4B and 4C**). These results suggest that both enrichment of importin α at the membrane and depletion of importin α at the membrane can both result in misregulation of cilia length. Thus, a balance between the levels of membrane-bound and cytoplasmic importin α must be maintained for proper cilia length control. These two populations of importin α may play distinct but equally critical roles in ciliary trafficking, both of which are regulated by the palmitoylation of importin α.

**Figure 4:**
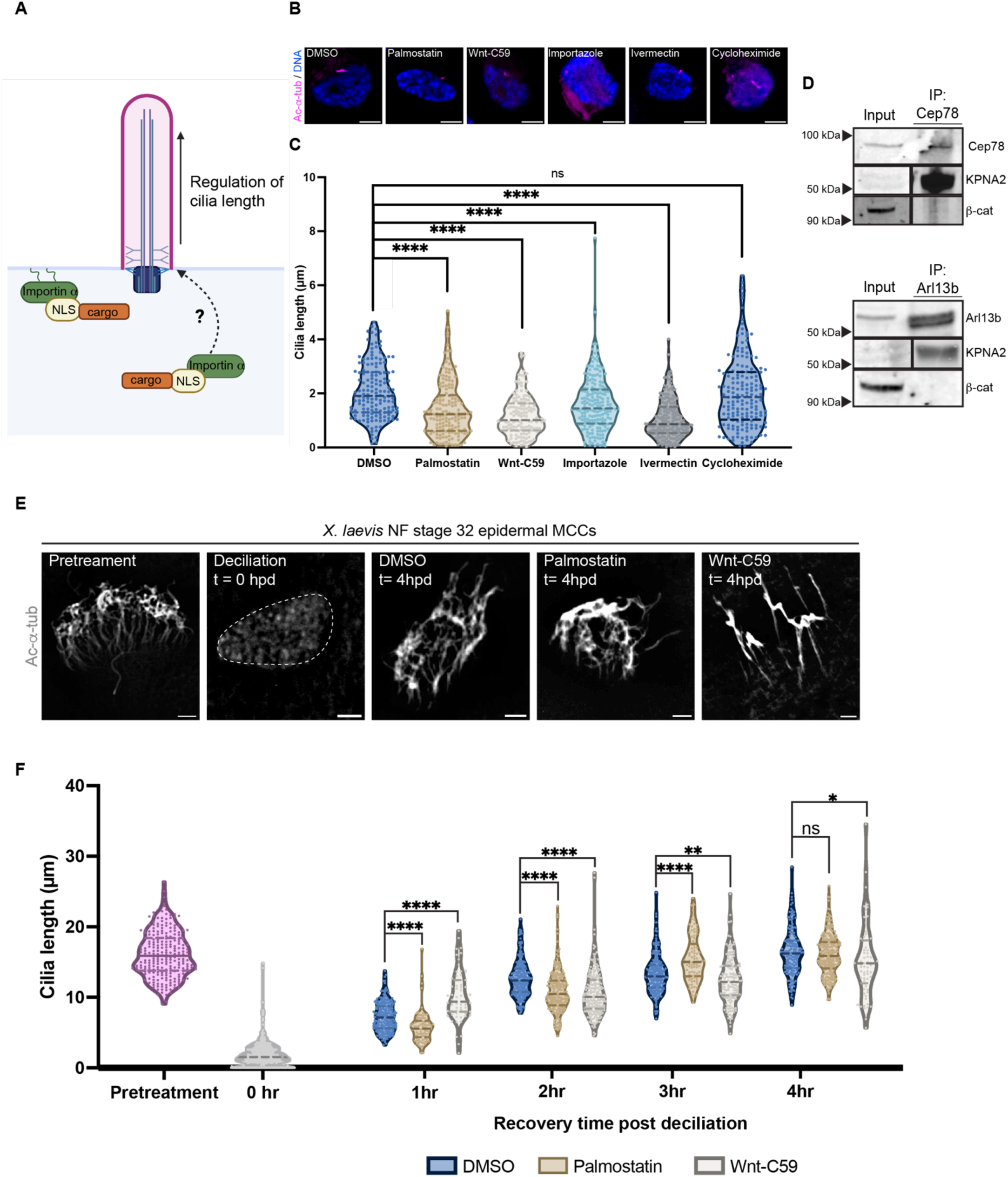
Cilia length is regulated by the balance of palmitoylated and unpalmitoylated (cytoplasmic) importin α. **A)** Schematic of hypothesized role for palmitoylated importin α and cytoplasmic importin α regulating proper cilia length. **B)** Deconvolved epifluorescent images of ciliated RPE-1 cells treated for 2 hours with drugs: DMSO, 10μM Wnt-C59, 50μM palmostatin, 40μM importazole, 25µM Ivermectin, or 50µM Cycloheximide, immunostained for acetylated-α-tubulin (magenta), and DNA (blue). Scale bar = 5µm**. C)** Quantification of cilia length (µm) of ciliated RPE-1 cells treated for 2hrs with drugs. ***P<0.001, ****P<0.0001 Student’s t-test, n=100 cells per treatment, 3 replicates. **D)** Immunoblot of RPE-1 cells immunoprecipitated for Cep78 or Arl13b and immunoblotted for each of the respective antibodies and KPNA2 (importin α) and β-catenin. **E)** *X. laevis* NF stage 32 epidermal MCCs were chemically deciliated and allowed to reciliate over 4 hours. Deconvolved epifluorescent images of *X. laevis* MCCs pretreatment, deciliated MCCs (t= 0 hours post deciliation, hpd) and regrown MCCs 4 hours post deciliation for MCCs recovered in DMSO, 100µM Wnt-C59 or 1mM palmostatin, immunostained for acetylated-α-tubulin (gray). Scale bars = 5µm. **F)** Quantification of cilia length (µm) before deciliation and 4hrs of recovery time post deciliation. * P<0.05, ** P< 0.01, **** P<0.0001, Student’s t-test, n= 90 cilia, n≥2 replicates per condition.

Next, we explored the significance of importin function in releasing and binding cargo proteins during steady state dynamics of cilia length maintenance. To achieve this, ciliated RPE-1 cells were treated with importazole, which prevents the release of cargo bound to importin α, which resulted in a reduction in cilia length (**Figures 4B and 4C**). To understand the requirement for importin α binding to cargo in regulating cilia length, ciliated RPE-1 cells were treated with the competitive inhibitor of importin α cargo binding, ivermectin^44^. Upon addition of ivermectin, cilia length was also decreased, suggesting a role for importin α binding of cargo as being necessary in regulating cilia length (**Figures 4B and 4C**). To eliminate the potential confounding effect of abrogating normal transcription during the experimental duration in maintaining ciliary length, a side-effect of inhibiting importin α-NLS binding/release, we utilized cycloheximide, an inhibitor of eukaryotic translation elongation^45^. Cycloheximide treatment of ciliated RPE-1 cells did not significantly alter cilia length, indicating that cilia can maintain their normal length without new protein synthesis during the experimental period. Overall these data suggest that there is a prerequisite transport system of importin α in both palmitoylated and cytoplasmic forms to bind and release cargo proteins, facilitating their trafficking into the cilium to maintain proper cilia length. Our data has demonstrated that importin α is required for both cilia formation and cilia length maintenance. However, the ciliary cargos that require importin α binding to facilitate these processes remains unknown. To address this, we conducted a bioinformatic screen to identify potential NLS-containing ciliary proteins that could potentially interact with importin α. We utilized three independently published NLS prediction tools; NucPred, cNLS Mapper, and seqNLS^46–49^ (**Figure S4**). From the bioinformatic screen, we identified two potential NLS-containing ciliary cargos involved in specifically regulating cilia length: Centrosomal protein (Cep78) and Arl13b. Cep78 is a centrosomal protein that has been identified to be critical for cilia length regulation through its interaction with the CP110 complex^50^. Arl13b is a small regulatory GTPase ciliary protein that localizes to the ciliary membrane and has been characterized to regulate cilia length^51^. After identifying Cep78 and Arl13b as cilia length regulators and containing NLS sequences, we performed co-immunoprecipitation analysis to determine if importin α can bind to these proteins. Immunoblot analysis revealed that importin α (KPNA2) directly interacts with both Cep78 and Arl13b (**Figure 4D**). After establishing the role of importin α in regulating primary cilia length in RPE-1 cells, we tested its role in MDCK-II cells and observed a similar trend. This suggests that importin α plays a role in regulating primary cilia length in cells that use either the intracellular or alternative ciliogenesis pathway. (**Figures S3B and S3C**).

### Importin α regulates cilia length in *X. laevis* epidermal MCCs

Next, we sought to investigate whether importin α also regulates cilia length in *X. laevis* epidermal MCCs. Another advantage of the *X. laevis* epidermal MCC model system is that in addition to studying steady state dynamics of cilia length regulation, the deciliation technique demonstrated in Figure 1G can be used to observe changes in cilia regrowth kinetics in real time. As previously mentioned, the deciliation process sheds cilia from MCCs, followed by cilia regrowth over a 4-hour time period^25^. This rapid regrowth period provides a valuable window to assess the impact of importin α on cilia length regulation. Therefore, after deciliation is performed, *X. laevis* embryos were recovered in a drug bath of either palmostatin or Wnt-C59 to determine the effect of inhibiting importin α membrane localization on cilia length regrowth kinetics during the regeneration period (**Figure 4E**). While cilia length varied over time between both treatments, at the final time point the cilia regrown in the Wnt-C59 recovery embryos were significantly shorter compared to control and palmostatin recovery embryos (**Figure 4F**). Additionally, while the cilia were shorter in the Wnt-C59 recovery embryos, they also contained fewer cilia within the epidermal MCC. These results support our previous data suggesting the dual role for importin α in regulating cilia formation and cilia length. Altogether, our data shows that importin α plays a role in cilia length regulation in both steady state dynamics and during cilia regrowth.

### Importin α binds to the NLS sequence of Cep164 to mediate ciliogenesis

The previously mentioned bioinformatics screen employed to identify putative NLS-containing ciliary proteins also identified the centrosomal protein, Cep164 (**Figure S4**) as likely to contain a bona-fide NLS sequence. Cep164 has been extensively studied as a protein which localizes to the distal appendages of the mother centriole and is essential for proper basal body docking, thereby facilitating cilia formation^52^. Given that Cep164 is predicted to contain a NLS sequence, we sought to determine if importin α could potentially bind Cep164 through its NLS sequence to mediate its localization for proper ciliogenesis. Initially, we performed immunofluorescent analysis and confirmed the co-localization of importin α and Cep164 in ciliated RPE-1 cells and deciliated *X. laevis* epidermal MCC for visualization at the MCC base (**Figures 5A, 5C and 5D**). To enhance the visualization of importin α co-localizing with Cep164 we performed U-ExM to increase resolution and found importin α localizing in a similar pattern to that of Cep164 and the distal appendages (**Figure 5B**). We next utilized the DuoLink Proximity Ligation Assay (PLA) in order to determine whether this co-localization occurs within a 40nm distance, indicative of a direct interaction. The PLA was performed on ciliated RPE-1 cells and revealed the binding of importin α and Cep164 at the cilium base (**Figures 5E and 5F**). To confirm that the binding of importin α and Cep164 is mediated by its NLS sequence, we generated a mutant version of Cep164 by alanine substitution of the positively charged lysine residues found within the predicted NLS sequence (Cep164 ∆NLS GFP) (**Figure 5G**). RPE-1 cells were transfected with Cep164 ∆NLS GFP and a wild-type construct^52^ (Cep164 WT GFP) and immunoblot analysis demonstrated that importin α binds to Cep164 via its NLS sequence (**Figures 5H and 5I**). To test whether importin α binding to Cep164 via its NLS sequence is required for proper ciliogenesis, we transfected RPE-1 cells with Cep164 WT GFP and Cep164 ∆NLS GFP plasmids, along with Arl13b-mCardinal for cilia visualization^52,53^. The overexpression of Cep164 ∆NLS GFP exhibited a decrease in ciliation, indicating a dominant-negative effect on ciliogenesis (**Figures 5J and 5K**). In summary, our data elucidates the role of importin α in mediating the transport of ciliary cargo proteins, such as Cep164, Cep78, and Arl13b for proper regulation of ciliogenesis and cilia length (**Figures 4,5**).

**Figure 5:**
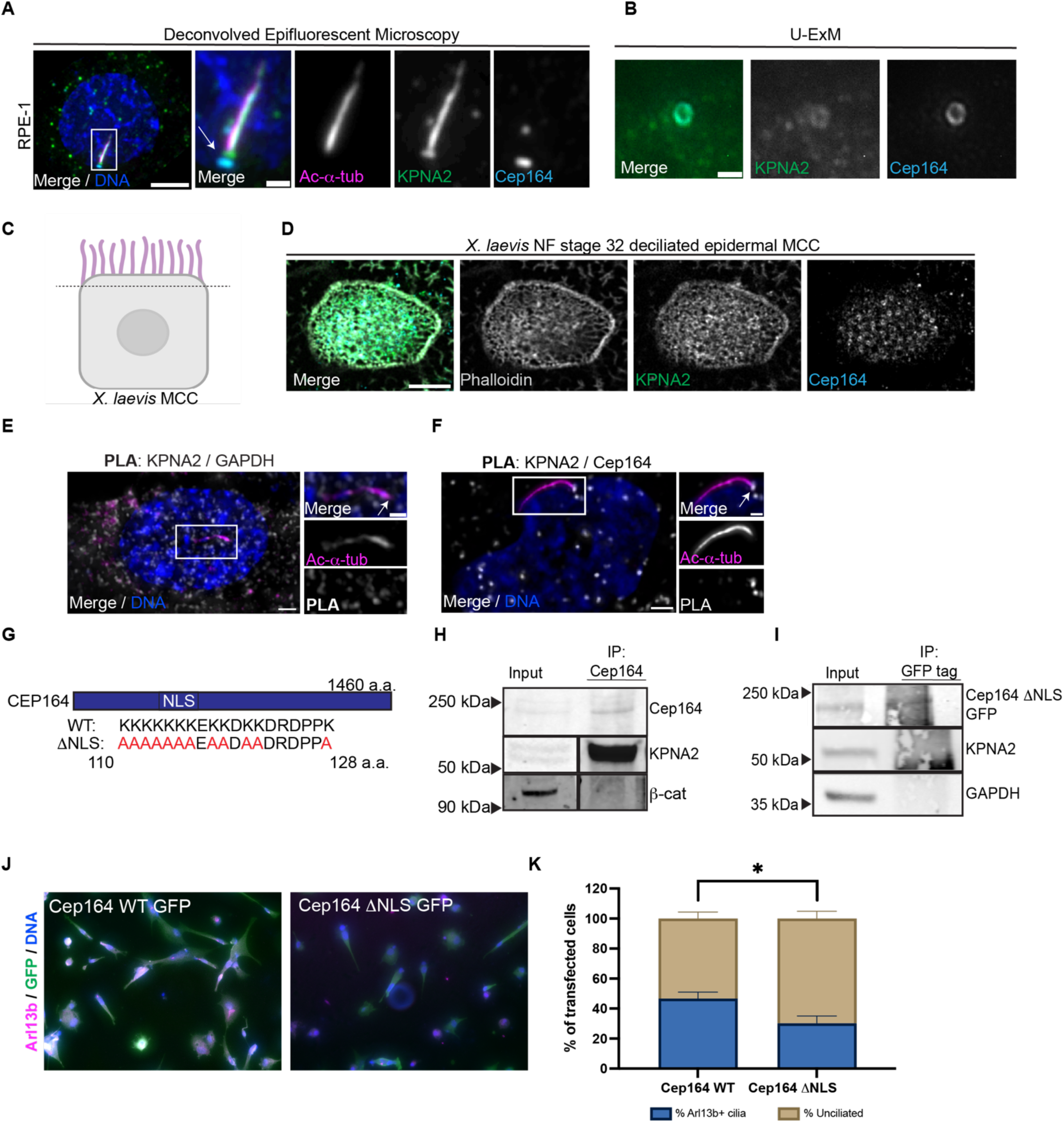
Cep164 ∆NLS cannot bind importin α and acts as a dominant negative mutant inhibiting ciliogenesis. **A)** Deconvolved epifluorescent image of a ciliated RPE-1 cell immunostained for acetylated-α-tubulin (magenta), KPNA2 (importin α) (green), distal appendage protein, Cep164 (cyan) and DNA (blue). Scale bars = 5µm (full image) and 1µm (inset images). **B)** U-ExM deconvolved epifluorescent image top-down view of a ciliated RPE-1 cell immunostained for KPNA2 (importin α) (green) and Cep164 (cyan). Scale bar = 1µm. **C)** Schematic of a deciliated *X. laevis* MCC, for a top-down view of importin α localization at MCC base. **D)** Airyscan confocal images of a *X. laevis* NF stage 32 deciliated epidermal MCC immunostained for phalloidin (gray), KPNA2 (importin α) (green) and Cep 164 (cyan). Scale bar = 5µm. **E)** Deconvolved epifluorescent image of a Proximity Ligation Assay (PLA) performed on ciliated RPE-1 cells, negative control with KPNA2 (importin α) and GAPDH (gray). Zoomed in image of lack of PLA spot at base of cilia (white arrow), acetylated-α-tubulin (magenta) and DNA (blue). **F)** Deconvolved epifluorescent image of PLA performed on RPE-1 cells for KPNA2 (importin α) and Cep164 (gray). Zoomed in image of PLA spot at cilium base (white arrow), acetylated-α-tubulin (magenta) and DNA (blue). Scale bars = 2µm (full image) and 1µm (inset images). **G)** Predicted nuclear localization signal (NLS) sequence within Cep164 spanning amino acid (AA) residues 110-128. Generated a mutant version of the Cep164 NLS (Cep164 **∆**NLS GFP) by replacement the lysines (K) with alanines (A). **H)** Immunoblot of RPE-1 cells immunoprecipitated for Cep164, immunoblotting for Cep164, KPNA2 (importin α), and β-catenin. **I)** RPE-1 cells transfected with 1µg Cep164 ΔNLS GFP and immunoprecipitated for GFP tag of construct. Immunoblotting for GFP, KPNA2 (importin α), and GAPDH. **J)** Epifluorescent images of RPE-1 cells transfected with 750ng of Arl13b-mCardinal and 250ng of either Cep164 WT GFP or Cep164 **∆**NLS GFP immunostained for plasmid with GFP (green), Arl13b (magenta), and DNA (blue). After transfection, RPE-1 cells were serum starved to induce ciliogenesis. Scale bars = 50µm. **K)** Quantification of % of Arl13b+ cilia and % of unciliated cells in transfected RPE-1 cells from Figure 5J. * P<0.05 Student’s t-test n=100 cells per treatment, 4 replicates. Mean ± SEM.

### Disruption of palmitoylated importin α contributes to a *X. laevis* kidney ciliopathy phenotype

Overall, this study has demonstrated the indispensable role of palmitoylated importin α in regulating ciliogenesis and cilia length through its interaction with different NLS-containing cilia cargo proteins. Building upon this, we aimed to investigate whether disrupting palmitoylated importin α in these processes could contribute to a ciliopathy phenotype. Ciliopathies can manifest in various organs and for this study we focused on the kidney due to numerous studies that have linked proper primary cilia function in the kidney with proper kidney morphology^3,4,53,54,55^. It is thought these progressive changes stem from signaling defects due to either the absence of cilia or aberrant cilia length^54–56^. As we’ve demonstrated altered cilia length in MDCK-II cells (kidney epithelial cells) by disrupting the palmitoylation of importin α (**Figure S3**), we sought to leverage the *X. laevis* model to explore the potential contributions of palmitoylated importin α in renal ciliopathy.

*X. laevis* are an ideal model to study kidney development (nephrogenesis) because its embryos contain one nephron, the functional unit of the kidney on each side of its body, while human kidneys normally contain 200,000 to over 2.5 million nephrons^57^. The genetic networks that regulate mammalian nephrogenesis and nephron morphology are analogous to the *X. laevis* pronephros, also known as the embryonic kidney which forms around NF stage 42 during *X. laevis* development^58,59^ (**Figure 6A**). The *X. laevis* embryonic kidney is comprised of tubular sections, such as the distal and proximal tubules which are known to filter waste products and reabsorb nutrients from the blood^59^. To elucidate the potential contributions of palmitoylated importin α in *X. laevis* nephrogenesis, embryos were treated with Wnt-C59, inhibiting importin α palmitoylation. Given the potential confounding effects of Wnt-C59 treatment on the Wnt pathway, embryos were also treated with okadaic acid (OA), a PP1 and PP2A phosphatase inhibitor. Previous studies have demonstrated that OA can activate the Wnt pathway downstream and independent of Wnt secretion, thereby rescuing the effects of PORCN inhibition on Wnt signaling^60,61^. Consequently, we hypothesized that treatment with Wnt-C59 and OA would inhibit importin α at the membrane via palmitoylation while preserving active Wnt signaling within the *X. laevis* embryo. Upon treatment of *X. laevis* embryos with Wnt-C59, OA or Wnt-C59 + OA, we observed significant kidney morphology defects, characterized by the loss of tubular structures and the development of enlarged regions (potentially cysts), reminiscent of those observed in kidney ciliopathies (**Figures 6B and 6C**). These results demonstrate that palmitoylation of importin α specifically is required for proper *X. laevis* nephrogenesis.

**Figure 6:**
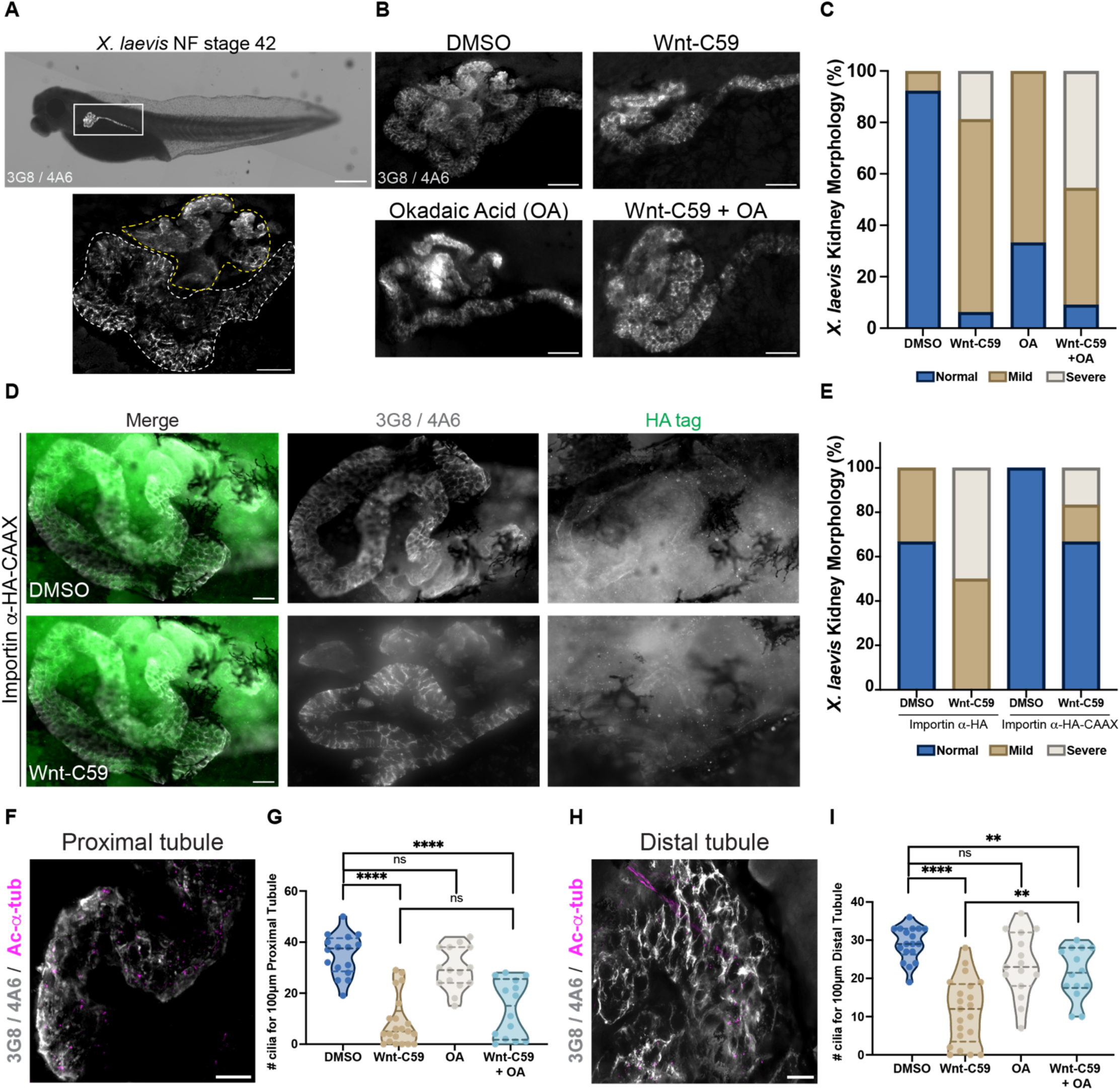
Disruption of importin α palmitoylation causes *X. laevis* renal ciliopathy. **A)** Deconvolved epifluorescent image of a *X. laevis* NF stage 42 embryo immunostained for the proximal tubules (3G8) and distal tubules (4A6) (gray), comprising of the embryonic kidney, outlined in the white box. Scale bar = 200µm. Below is a deconvolved epifluorescent image of a *X. laevis* NF stage 42 embryonic kidney (gray) with the proximal and distal tubule outlined in yellow and white dashed lines respectively. Scale bar = 100µm. **B)** Deconvolved epifluorescent images of *X. laevis* embryonic kidneys (gray) drug treated with DMSO, 100µM Wnt-C59, 10nM Okadaic Acid (OA) or 100µM Wnt-C59 + 10nM OA until embryos reached NF stage 42. Scale bar = 100µm. **C)** Quantification of % *X. laevis* embryonic kidneys with normal, mild or severe kidney morphology for all drug treatments. Threshold of determining kidney morphology phenotypes is outlined in the Material and Methods section. n≥14 tadpoles per condition. **D)** Deconvolved epifluorescent image of a *X. laevis* embryos microinjected at the 4-cell or 8-cell stage with 60pg of importin α-HA-mCH-CAAX targeted to the kidney. After microinjection, embryos were treated with either DMSO or 100 µM Wnt-C59 until NF stage 42, immunostained for embryonic kidney (gray) and HA tag (green). Scale bar = 20µm. **E)** Quantification of % *X. laevis* embryonic kidneys with normal, mild or severe kidney morphology for all drug treatments. n≥6 tadpoles per condition. **F)** Deconvolved epifluorescent image of magnified proximal tubule (gray) immunostained for acetylated-α-tubulin (magenta). Scale bar = 20µm. **G)** Quantification of # of cilia for 100µm of proximal tubule for all drug treatments. ****P<0.0001. **H)** Deconvolved epifluorescent image of magnified distal tubule (gray) immunostained for acetylated-α-tubulin (magenta). Scale bar = 20µm. **I)** Quantification of # of cilia for 100µm of distal tubule for all drug treatments. Student’s t-test, ** P< 0.01, **** P<0.0001, n≥15 tadpoles per condition.

To investigate if forced membrane localization of importin α via the CAAX motif is sufficient to rescue the kidney morphology defects observed with Wnt-C59 treatment, we microinjected embryos with importin α-mCH-HA or importin α-mCH-HA-CAAX plasmids targeted to the blastomere fate-mapped to give rise to the embryonic kidney and treated injected embryos with DMSO or Wnt-C59 until NF stage 42. While neither importin α-HA nor importin α-mCH-HA-CAAX expression alone altered kidney morphology (**Figures S5A-S5C**), remarkably, we observed a rescue in kidney morphology in our importin α-mCH-HA-CAAX + Wnt-C59 treatment embryo, supporting our hypothesis that membrane-bound importin α is required for proper nephrogenesis (**Figures 6D and 6E**). After characterizing the global effect of inhibiting importin α palmitoylation, we next sought to investigate how renal primary cilia in the distal and proximal tubule segments of the *X. laevis* kidney are affected. In the proximal tubule region, we observed a lack of primary cilia rescue in our Wnt-C59 +OA treatment, suggesting that under these conditions, importin α palmitoylation is essential for proper primary cilia formation (**Figures 6F and 6G**). Conversely, we did not observe this effect in the distal tubule region, as there was a partial rescue of primary cilia present in this treatment (**Figures 6H and 6I**). Our data suggest while palmitoylated importin α may not be necessary in both segments of the *X. laevis* kidney, there is still a certain requirement for palmitoylated importin α in regulating renal primary ciliogenesis for overall proper nephrogenesis.

## Discussion

Our study offers a previously unrecognized function for palmitoylated and non-palmitoylated importin α in regulating ciliogenesis and cilia length in both human and *X. laevis* renal primary cilia and *X. laevis* epidermal multicilia. Previous research has established the localization of RanGTP, nuclear pore proteins, and importin β as localizing to the cilia and their essential role in ciliogenesis^10–13^. We have identified a pathway that integrates these findings by characterizing the localization of palmitoylated and non-palmitoylated importin α at the cilia base and lumen, respectively, as well as multiple NLS-containing cargos required for ciliogenesis and cilia maintenance (**Figure 7**). This study builds upon previous findings that demonstrate the role of palmitoylated importin α in regulating membrane-associated processes, including nuclear and spindle size scaling, mitotic spindle orientation, and now ciliogenesis^14,15^. Furthermore, our results suggest that palmitoylated importin α is crucial during nephrogenesis and is essential for renal primary cilia formation in the tubule segments of the *X. laevis* embryonic kidney (**Figure 7**).

**Figure 7:**
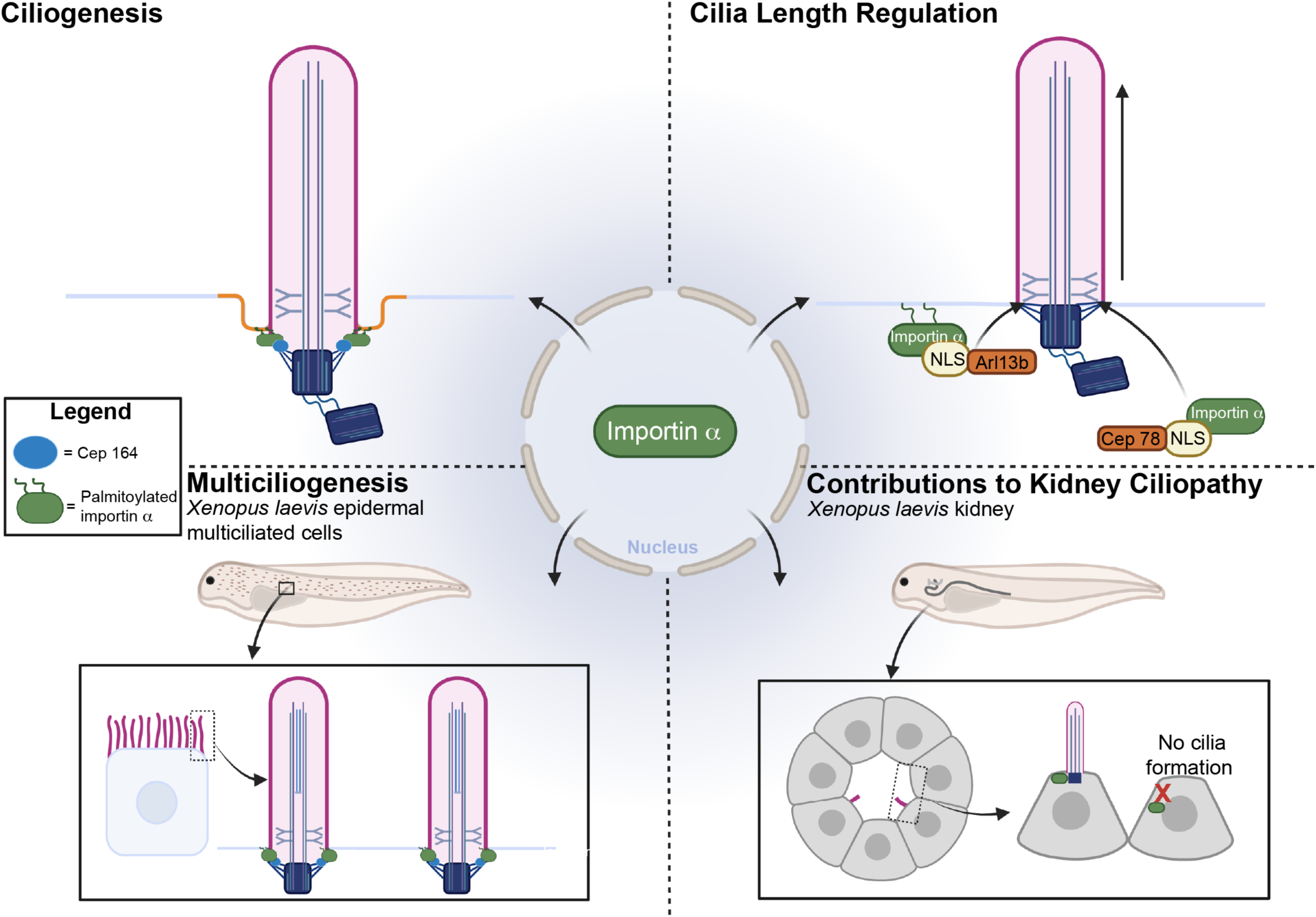
Importin α regulates ciliogenesis and cilia length maintenance in a palmitoylation dependent manner and is essential for proper kidney development. Importin α possesses a novel function in the cell apart from its classic role in nuclear transport. This newly described function includes regulating ciliogenesis and cilia length maintenance in both primary and multiciliated cells. In addition, palmitoylated importin α is required for nephrogenesis in *X. laevis* embryos.

We have characterized the localization of importin α at ciliary structures through confocal microscopy and U-ExM (**Figure 1**). However, this localization raises the question of how and where palmitoylated importin α associates with the primary cilia membrane specifically. Current research in the field has identified the basal body distal appendage protein, Cep164, as localizing to the ciliary pocket in one of the initial steps of basal body docking prior to ciliogenesis^5,62^. Notably, cholesterol and lipids normally found in lipid raft microdomains have been demonstrated to be enriched at the ciliary pocket^27,63^. Given that palmitoylated proteins are known to associate with lipids rafts^64,65^, it is an intriguing possibility that palmitoylated importin α might localize to the ciliary pocket, where it could interact with Cep164 to facilitate basal body docking (**Figure 5**). This hypothesis is further supported by our findings that Cep164 directly associates with importin α at the base of the primary cilium via PLA (**Figures 5E and 5F**), and that Cep164 and importin α binding is mediated via the Cep164 NLS sequence as demonstrated via IP (**Figures 5G-5I**). This hypothesis may also explain the reduction in cilia observed when Cep164 ∆NLS GFP is overexpressed in RPE-1 cells, by functioning as a dominant-negative mutant. In this case, the intact C-terminus of Cep164 would still enable distal appendage binding, while the mutated NLS would prevent Cep164 binding of importin α at the ciliary pocket, thereby hindering docking and ciliogenesis^62^ (**Figures 5J and 5K**).

We have demonstrated that palmitoylated importin α is essential for proper ciliogenesis by disrupting its palmitoylation status with pharmacological inhibitors and siRNA knockdown of PORCN. However it remains unclear how importin α is initially associated with the nascent cilia (**Figure 2**). During the beginning of the intracellular ciliogenesis pathway, the mother centriole acquires Golgi-derived vesicles that dock onto the distal appendages mediated through Cep164. These vesicles fuse together to allow for the cilium to begin growing internally and then migrate apically to invaginate at the plasma membrane with the cilium continuing to grow. Previous studies exploring this process have demonstrated that the Golgi-derived vesicles recruited to the distal appendages contain ciliary proteins that range from Rab GTPases to various centrosomal proteins required to initiate ciliogenesis ^5,35,52,62^. In addition, previous work on the early stages of the alternative ciliogenesis pathway have linked shared ciliary proteins from the intracellular ciliogenesis pathway such as Rab GTPases and intraflagellar transport proteins known to congregate at the plasma membrane for cilia formation^67,68^. Of particular interest, it has been shown that importin α can also localize to the Golgi to undergo its palmitoylation modification for subsequent membrane localization^63,69,70^. Our data raises the intriguing possibility that palmitoylated importin α may be required to bind Cep164 within Golgi-derived vesicles before being transported to the growing cilium during the intracellular ciliogenesis pathway, in addition to its role in the alternative pathway involving basal body docking to the plasma membrane.

Our bioinformatics screen for NLS-containing ciliary proteins has identified Arl13b, a cilia membrane-bound protein, and Cep78, a non membrane-bound protein, both of which are required for regulating cilia length. While we demonstrated importin α can bind both proteins via IP (**Figure 4**), it is unclear which is binding either membrane-bound or non membrane-bound importin α. As we have shown the importance of maintaining a balance of importin α at the membrane and the cytoplasm to maintain proper cilia length, it will be important to elucidate which fraction of importin α is binding to which ciliary cargos. Additionally, studies have observed that Arl13b not only regulates cilia length but is also an essential factor for protein trafficking and the execution of Sonic hedgehog signaling (Shh)^51^. Shh signaling in cilia has been studied extensively and has been shown to be critical for the development and maintenance of various organs^51,66^. Further investigation promises to elucidate if the trafficking of Arl13b by importin α has the potential to impact Shh signaling and will reveal new connections between membrane-bound importin α in regulating downstream signaling events.

Taken together, the results of this study provide foundational insight into the understanding of the molecular mechanisms regulating ciliogenesis, protein transport to the cilia base, and cilia length maintenance. We demonstrate a previously unrecognized regulatory role for the nuclear transport adapter, importin α, in cilia formation through the binding of various NLS-containing ciliary proteins. These results introduce a new pathway which highlights an unexplored area of research in the current field linking importins to ciliogenesis and cilia length maintenance. These findings have profound implications for our understanding of the etiology of ciliopathies and the development of preventative and therapeutic strategies.

## Resource Availability Lead Contact

Requests for further information should be directed to and will be fulfilled by the lead contact, Christopher W. Brownlee.

## Materials Availability

Requests for resources and reagents should be directed to the lead contact. All plasmids generated in this study are being deposited to Addgene.

## Data and Code Availability

- Microscopy data reported in this study will be shared by the lead contact upon request
- No original code was generated in this study
- Any additional information required to reanalyze the data reported in this paper is available from the lead contact upon request

## Acknowledgements

We thank Patrick Sutton, Kathryn Malone, and Melanie Garcia for their constructive feedback on this work, engaging discussions, and edits during the writing of this manuscript. We thank Ken-Ichi Takemaru, Holly Colognato, Gerald Thomsen, and Helen Rankin Willsey for their insights and suggestions on experimental approach. We thank the Stony Brook University Central Microscopy Imaging Center (CMIC) and Guowei Tian for assistance in imaging with Zeiss LSM 980 Airsycan 2 NLO Two-Photon Confocal Microscope. Work was supported by National Institutes of Health grant 1R35GM147569-01 (C.W.B).

## Author Contributions

Conceptualization, C.W.B; Methodology, N.M. and C.W.B.; Investigation, N.M. and P.J.S.; Writing-Original Draft Preparation, N.M. and C.W.B.; Writing-Review & Editing, N.M., P.J.S., and C.W.B.; Funding Acquisition, C.W.B.; Resources, C.W.B.; Supervision, C.W.B.

## Declarations of Interests

The authors declare no competing interests.

## STAR★METHODS

### KEY RESOURCES

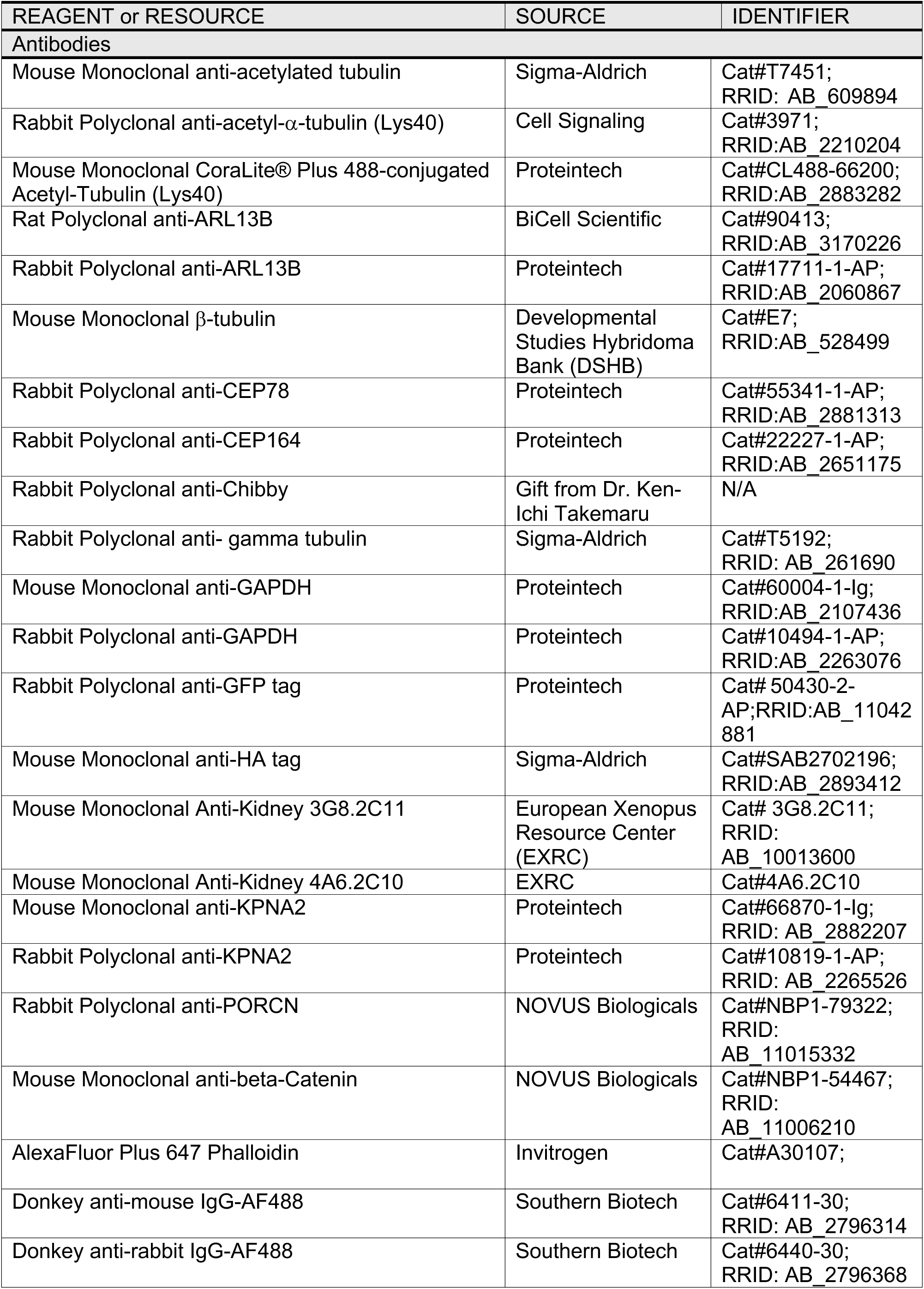
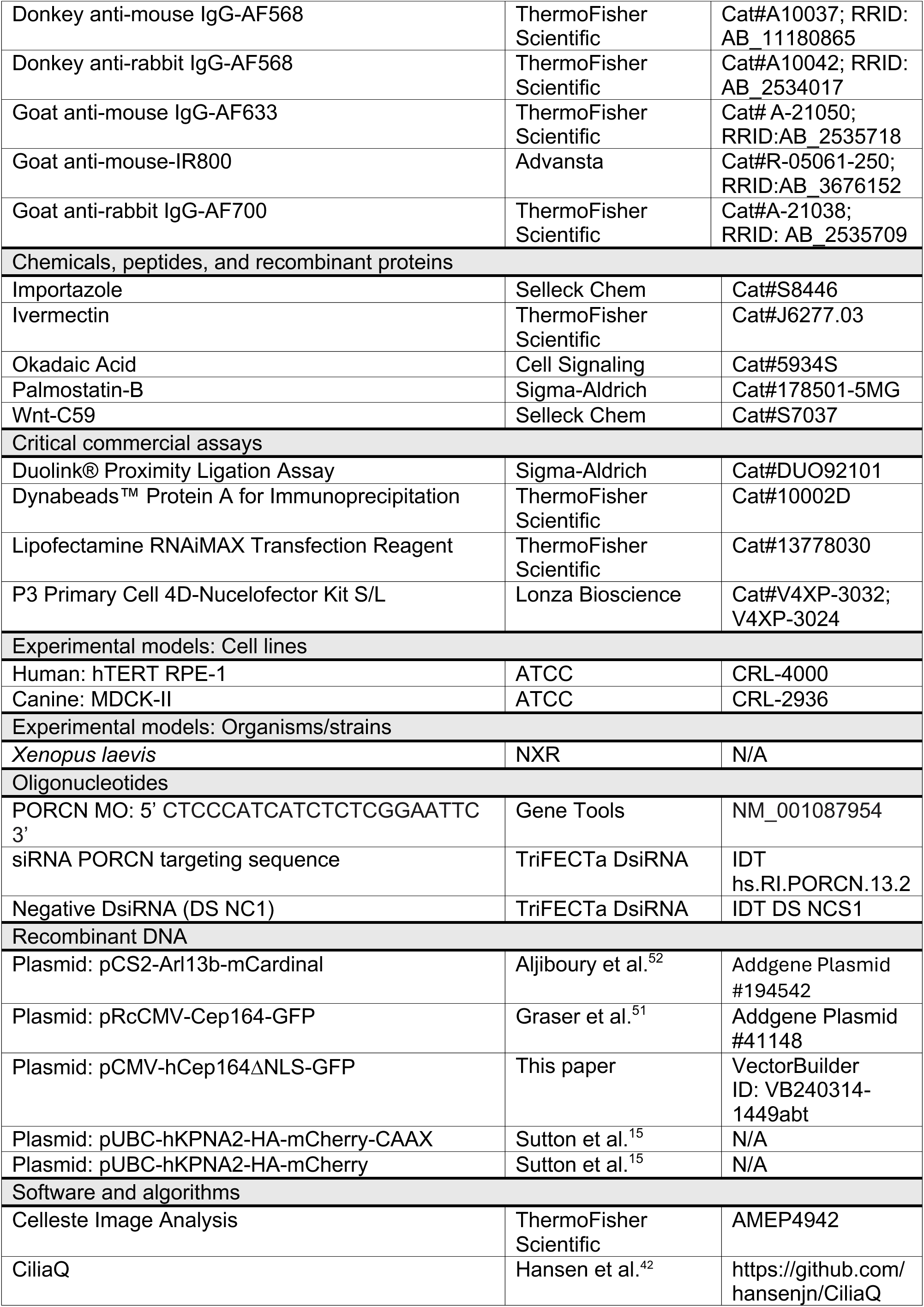
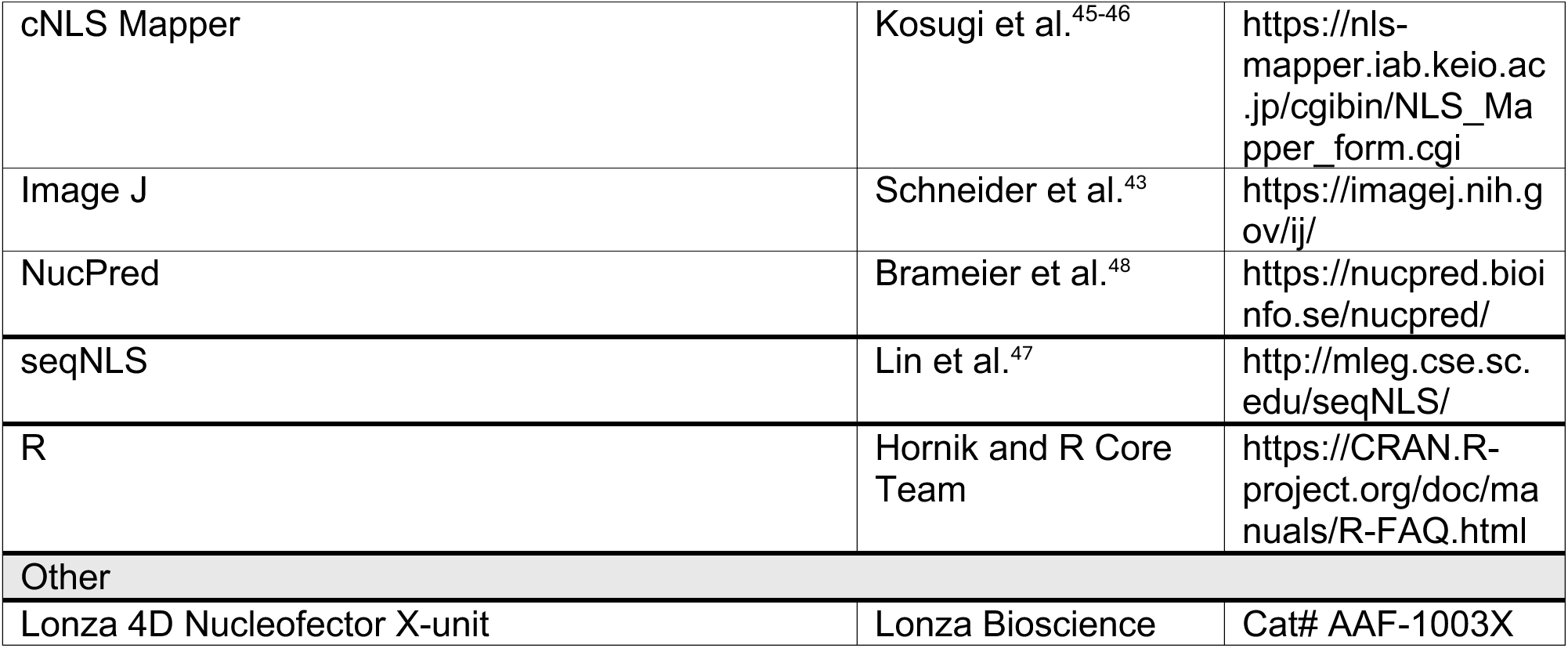

### METHOD DETAILS

#### Cell culture, drug treatments, and RNAi

hTERT-RPE-1 (RPE-1) and canine MDCK-II cells were obtained from ATCC. RPE-1 cells were grown in DMEM/F12 media supplemented with 10% v/v fetal bovine serum (FBS; VWR) and 1% v/v antibiotic-antimycotic (Anti-Anti; ThermoFisher Scientific). MDCK-II cells were grown in EMEM media supplemented with 10% v/v FBS and 1% v/v Anti-Anti. Cell lines were grown at 37°C and 5% CO_2_ in a humidified chamber. To induce ciliogenesis RPE-1 cells were serum starved for 48 hours and MDCK-II cells were grown in complete media for 120 hours. For ciliogenesis analysis, MDCK-II cells were cultured on transwell 0.4µm pore size inserts (Fisher Scientific). Cells were routinely checked for mycoplasma using the Mycoplasma Detection kit (Southern Biotechnology). Pharmacological drug treatments for palmitoylation and importin function were added at concentrations listed in figure legends in DMEM F12 or EMEM serum free media 24 hours prior to ciliogenesis for RPE-1 and MDCK-II cells respectively. RNAi experiments on RPE-1 cells were performed using the TriFECTa DsiRNA kit. The oligonucleotides for PORCN and a negative control were transfected at an amount of 30 pmol per 6 well plate using the Lipofectamine RNAiMAX transfection reagent 3000 in accordance with the supplier’s recommendations. RPE-1 cells were knockdown with siRNA PORCN for 24hrs prior to serum starvation of 48hrs to induce ciliogenesis. Samples were either fixed for immunostaining of prepared for cell lysis for immunoblotting as described below.

#### Ultra-expansion microscopy (U-ExM)

RPE-1 cells were grown on 12mm fibronectin coated coverslips using fibronectin bovine plasma (ThermoFisher Scientific #33010018) at 5µg/mL. RPE-1 cells were serum starved for 48hr to induce ciliogenesis. Once ciliated, RPE-1 cells were fixed with 4% PFA in PBS for 2 hours at room temperature and then proceeded to the U-ExM protocol as defined by Laporte et al. Primary antibodies used for U-ExM; mouse anti-acetylated-α-tubulin (1:250, Sigma-Aldrich), rabbit anti-gamma-tubulin (1:250, Sigma-Aldrich), mouse anti-KPNA2 (1:100, Proteintech), and rabbit anti-Cep164 (1:200, Proteintech), Hoechst (1:1000, Cayman Chemical). To image expanded gels, poly-d-lysine (0.1mg/mL) coated slides were used for mounting and imaged on the EVOS M7000 epifluorescent microscope.

#### Animal models

All animals were maintained in accordance with standards established by the Division of Laboratory Animal Resources at Stony Brook University.

#### *Xenopus laevis* (*X. laevis*) in-vitro fertilization

X. laevis adult females were induced to lay eggs by a priming injection of 100 U pregnant mare serum gonadotropin (PMSG) at least 48 hours before use and a boosting injection of 500 U of human chorionic gonadotropin (hCG) 16 hours before use. Following the hCG injection, adult female *X. laevis* were placed in a 2L water bath of 1X MMR (100 mM NaCl, 2 mM KCl, 1 mM MgCl2, 2 mM CaCl2, 0.1 mM EDTA, 5 mM HEPES pH 7.8) overnight at 17°C. Approximately 16 hours following hCG injection, fresh eggs were collected by squeezing eggs from ovulating frogs into a 10cm plastic petri dish. To fertilize eggs, a sperm solution made from ¼ of a male frog testis was placed in 1mL 1X MR (100 mM NaCl, 1.8 mM KCl, 2.0 mM CaCl2, 1.0 MgCl2, 5.0 mM HEPES-NaOH, pH 7.6) and homogenized using scissors and a pestle. 1mL of sperm solution was added dropwise to the freshly squeezed eggs and the dish was swirled to form a monolayer of eggs and incubated for 3 mins. Dishes were flooded with milli-Q water and incubated for an additional 10 mins. Eggs were dejellied with a 2% cysteine solution for 6 minutes with occasional swirling and washed 5 times with 1/3 MR. Fertilized eggs were incubated at 23°C until the appropriate developmental stage as described by Nieuwkoop and Faber (NF) 1994. *X. laevis* embryos treated with DMSO, 100µM Wnt-C59, 1mM Palmostatin or 10nM Okadaic Acid in fresh 1/3 MR and drug treated until the developmental stage indicated throughout the text and figures when applicable.

#### Immunofluorescence staining

##### Cell culture

RPE-1 cells were grown on 12mm fibronectin coated coverslips. Polarized MDCK-II cells were grown on 12mm 0.4µm pore size transwell filters and once ciliated the membrane was cut out and fixed in 4% PFA for 10 minutes at RT. Following fixation, fixed cells were permeabilized with PBST (0.2% Triton X-100 in PBS, Sigma-Aldrich 9036-19-5) 3 x 5 min and blocked for 1hr at RT in Bovine Serum Albumin (BSA) -PBST (3% BSA in PBST). Primary antibodies used were: mouse anti-acetylated-α-tubulin (1:1000, Sigma-Aldrich), mouse anti-KPNA2 (1:250, Proteintech), rabbit anti-Chibby (1:500, Gift from Dr. Ken-Ichi Takemaru), mouse anti-HA tag (1:250, Sigma-Aldrich), rabbit anti-GFP tag (1:250, Proteintech), and rabbit anti-Cep164 (1:250, Proteintech). Following primary antibody incubation for 1hr at RT, coverslips were washed 3 x 5min with PBST and then incubated for 30min with secondary antibodies. Secondary antibodies used were: Donkey anti-mouse and anti-rabbit IgG-AF488 (Southern Biotech), Donkey anti-mouse and anti-rabbit IgG-AF568 and Donkey anti-mouse IgG-AF633 (ThermoFisher Scientific) and all used at 1:1000 dilution. For Figures 1B and 5A, following incubation with secondary antibodies, samples were incubated with CoraLite® Plus 488-conjugated Acetyl-Tubulin (Lys40) Monoclonal antibody at 1:200 for 2hr. Finally, coverslips were washed 3 x 5 min with PBST and mounted with ProLong Diamond Antifade Mountant (ThermoFisher #P36961) and imaged on EVOS M7000 microscope. RPE-1 cells (Figures 2-3) were imaged with an Olympus 60x, 0.7 NA oil objective. For localization of importin α and Cep164 in RPE-1 cells (Figure 5), they were imaged on EVOS M7000 epifluorescent microscope.

##### X. laevis

Whole *X. laevis* embryos at the appropriate stages indicated in figure legends were fixed in 4% PFA in PBS for 2h at RT or overnight at 4℃, followed by 3 x 5 min with PBST (0.2% Triton X-100 in PBS). After permeabilization embryos were blocked in BSA-PBST (3% BSA in 0.2% Triton X-100 in PBS) for 1h at RT followed by overnight primary antibody incubation at 4°C with rotation. Primary antibodies used were: rabbit anti-acetylated-α-tubulin (1:500, Cell Signaling), mouse anti-KPNA2 (1:250, Proteintech), mouse anti-HA tag (1:250, Sigma-Aldrich), Cep164 (1:250, Proteintech), Phalloidin (1:100, ThermoFisher Scientific), mouse proximal kidney tubules 3G8 (1:30; EXRC) and distal kidney tubules 4A6 (1:5, EXRC). Primary antibodies were washed out 3 x 1hr with PBST and embryos were incubated overnight at 4°C with secondary antibodies in PBST. Secondary antibodies used were: Donkey anti-mouse and anti-rabbit IgG-AF488 (Southern Biotech), Donkey anti-mouse and anti-rabbit IgG-AF568 and Donkey anti-mouse IgG-AF633 (ThermoFisher Scientific) and all used at 1:500. Secondary antibodies were washes out 3 x 1hr and mounted with ProLong Diamond antifade mountant (ThermoFisher Scientific) and imaged on EVOS M7000 or Zeiss 980 confocal microscope.

#### Ciliation quantification

For quantification of changes in ciliogenesis, or percentage of ciliated cells was performed by counting the number of ciliated cells vs. unciliated cells using the point tool in ImageJ (v2.0.0). Quantification was plotted with GraphPad Prism 10 and number of samples are indicated in figure legends. For siRNA PORCN treated RPE-1 cells, the stubby cilia phenotype was determined by a ciliated cell possessing a cilium that appeared punctate.

#### *X. laevis* deciliation and motile cilia isolation

*X. laevis* NF stage 30 embryos were subjected to the deciliation procedure as described previously by Werner and Mitchell, 2013. Briefly *X.laevis* embryos were deciliated with 75mM CaCl_2_ and 0.02% NP-40 in 1/3 MR for 2min or until cilia movement stopped. Deciliated embryos were then fixed immediately in 4% PFA and followed immunostaining protocol above, or deciliated embryos recovered in a drug bath of either DMSO, 100µM Wnt-C59 or 1mM palmostatin in 1/3x MR, with subsequent fixation and immunostaining after each hour for a total of 4 hours post deciliation for cilia regrowth. Motile cilia were isolated and prepared for immunoblotting as previously described by Seidl, C., et al., 2023.

##### Quantification of reversible deciliation

Quantification of cilia regrowth after deciliation was performed using the Celleste Image Analysis Software (ThermoFisher Scientific). MCCs in each image were first selected as ROIs using the Celleste ‘select’ tool. Next within each MCC/ROI, 3 clear individual cilia (from cilia base to tip) were traced using the direct measurement polyline tool. Cilia length was measured in microns (µm) and plotted with GraphPad Prism 10. Number of samples are indicated in figure legends.

#### *X. laevis* immunoblotting

Isolated cilia and whole *X.laevis* embryo (1 embryo) were collected as described above and mixed 1:1 with 2X Laemmli buffer, boiled for 10min at 70℃. and 5 min at 95℃. respectively. Samples were loaded for SDS-PAGE in a 7.5% Tris-glycine gel and then transferred on to a nitrocellulose membrane. The membrane was then washed 3 x 5min with PBST (0.1% Tween-20 and 0.01% SDS in 1x PBS) and incubated with 5% milk blocking buffer for 1 hour at RT, followed by overnight primary antibody incubation at 4°C. Primary antibodies used were: rabbit anti-acetylated-α-tubulin (Cell Signaling), mouse and rabbit anti-KPNA2 (Proteintech), mouse and rabbit anti-GAPDH (Proteintech), and mouse anti-β-tubulin (DSHB) in blocking buffer all diluted 1:1000. Next the membrane was washed 3 x 5min with PBST (0.1% Tween-20 and 0.01% SDS in 1x PBS) and incubated with fluorescent secondary antibodies: IR-800 (1:10,000, Advansta) and IgG-AF700 (1:10,000, ThermoFisher Scientific) for 1 hour at RT, washed 3 x 5min with PBST and then scanned with an Odyssey Infrared Imaging System (LI-COR Biosciences).

#### *X. laevis* plasmid and morpholino microinjections

For epidermal MCC experiments, 30pg of importin α-mCH-HA or importin α-mCH-HA-CAAX, along with 5ng/nL of dextran as a tracer were injected targeted for epidermis at the 4-cell stage. For embryonic kidney experiments, 60pg of importin α-mCH-HA or importin α-mCH-HA-CAAX, along with 5ng/nL of dextran as a tracer at either the 4- or 8-cell stage targeted for the kidney. For morpholino experiments, PORCN morpholino was purchased from Gene Tools. *X. laevis* embryos were injected with 40ng of PORCN morpholino into one blastomere at the 2-cell stage, with the uninjected blastomere serving as an internal control . Morpholino sequence is: PORCN, (5’ CTCCCATCATCTCTCGGAATTC 3’). For PORCN morpholino injections, the injected half of the embryo was determined by observations in changes in pigment on the epithelium as the morpholino was target to the epidermis. All microinjected embryos were placed in 2.5% ficoll in 1/3MR with gentamycin for 4hrs, after which were then moved into a new dish of 1/3MR and incubated at 23°C. Drug treated and microinjected *X. laevis* embryos were treated with DMSO, 100µM Wnt-C59, or 10nM Okadaic Acid at NF stage 18-19 and grown until the appropriate developmental stage indicated in figure legends for fixation and staining.

#### *X. laevis* epidermal MCCs quantification

##### MCCs/Area

Quantification of the number of epidermal MCCs present on drug treated (DMSO, Wnt-C59 and palmostatin) *X. laevis* embryos was determined by an image taken with a 20x objective with an area of approximately 500 µm x 380 µm. Within this field the number of MCCs were counted with the point tool in Image J (v2.0.0). Number of samples are indicated in figure legends.

##### MCC cilia mask

For quantification of the integrated fluorescence density of the cilia within a MCC, the acetylated-α-tubulin channel was used for threshold “mean” segmentation in ImageJ (v2.0.0). The segmented cilia were selected for measuring the integrated density in arbitrary units and plotted with GraphPad Prism 10. Number of samples are indicated in figure legends.

#### *X. laevis* kidney morphology and cilia quantification

##### Kidney morphology

For quantification of *X. laevis* kidney morphology, the number of turns in the embryonic kidney, proximal and distal tubules, were counted with the point tool in ImageJ (v2.0.0). To categorize the number of turns into normal, mild or severe morphology, we established a threshold limit in which a kidney with 8 or more tubule turns are classified as normal, kidneys with less than 8 but greater than 4 turns were classified as mild and kidneys with less than 4 tubule turns were classified as severe. Morphology quantification was plotted with GraphPad Prism 10 and number of samples are indicated in figure legends.

##### Proximal and distal tubule cilia

For quantification of the number of cilia in the proximal and distal tubule regions, 100µm region of interest (ROIs) were made using ImageJ (v2.0.0). Within each ROI, the cilia were counted using the ImageJ point tool. Cilia quantification in the proximal and distal tubule were plotted with GraphPad Prism 10 and number of samples are indicated in figure legends.

#### Plasmid construct development

A mutated version of the Cep164 NLS, spanning amino acid resides 110-128, was generated by VectorBuilder. Mutation of the NLS was performed by replacing the lysine residues in the NLS to alanines, to preserve protein structure folding dynamics. This construct has the vector identifier #VB240314-1449abt.

#### Plasmid Transfection

RPE-1 cells were transfected via nucleofection LONZA P3 cell line 4D-Nucleofector kit (Catalog #V4XP-3032) following recommended protocol. Following nucleofection, RPE-1 cells were incubated in serum free media to induce ciliogenesis for a total of 48hrs. During this experimental time, 24hrs prior to ciliogenesis, RPE-1 cells were treated with either DMSO or 10µM Wnt-C59. After which, RPE-1 cells were fixed and immunostained as described above.

#### Cilia length quantification

RPE-1 cilia length was measured using the CiliaQ ImageJ plugins, CiliaQ Preparator (v0.1.2), CiliaQ Editor (v0.03) and CiliaQ (v0.1.7). The CiliaQ plugin was used as previously described by Hansen J.N, et al., 2021.

#### Immunoprecipitation and immunoblotting

Immunoprecipitation was performed according to manufacture protocol (ThermoFisher Scientific). Cell lysates were generated for immunoprecipitation experiments using a non-denaturing lysis buffer (20 mMTris HCl pH 8, 137 mM NaCl, 1% Triton X-100, 2 mM EDTA). supplemented with 10µg/mL each of leupeptin, pepstatin and chymostatin (LPC) protease inhibitors. Cell lysates were added 1:1 with 2X Laemmli sample buffer and subjected to western blot analysis. Antibodies used for immunoprecipitations and immunoblotting include: rabbit anti-Cep78, rabbit anti-ARL13B, rabbit anti-Cep164, rabbit anti-GFP tag, and mouse and rabbit anti-KPNA2 (all from Proteintech). Cell lysates were loaded on SDS-PAGE in a 7.5% Tris-glycine gel and proceeded through the protocol as described above. All antibodies used for immunoblotting were used at 1:1000. Blots were scanned with an Odyssey Infrared Imaging System (LI-COR Biosciences).

#### Proximity Ligation Assay (PLA)

PLA was performed using Duolink PLA fluorescence protocol (Sigma-Aldrich). RPE-1 cells were serum starved to induce ciliogenesis and fixed with 4% PFA in 1x PBS for 10 min. After fixation, samples were permeabilized with PBST for 15 min followed by blocking with the Duolink blocking solution for 1hr at 37℃. Next, samples were incubated with primary antibodies diluted in the Duolink Antibody Diluent for 1h at 37℃ and washed twice with Wash buffer A for 5 min each. Samples were then incubated with the Duolink Antibody Diluent containing mouse PLA^minus^ and rabbit PLA^plus^ probes for 1hr at 37°C followed by the ligation reaction for 30 min at 37℃. Next samples were incubated with the amplification reaction for 100 min at 37°C with the Duolink Fluorescent Detection Reagent Red. For cilia labeling, samples were then incubated for 2hrs at 37°C with CoraLite® 488-conjugated Acetyl-tubulin diluted in Duolink Antibody Diluent. Lastly, samples were washed with Wash buffer B and then mounted with the Duolink In Situ Mounting Media with DAPI for imaging on EVOS M7000 epifluorescent microscope with an Olympus Super Apo 100x, 1.4 NA oil objective.

#### NLS Bioinformatics Screen

NucPred (Brameier et al., 2007) was used to determine which cilia proteins contain potential NLS sequences with a threshold score of >0.63 (specificity > 71% and accuracy >53%). cNLS Mapper (Kosugi et al., 2009) was used to determine which cilia proteins contain potential NLS sequences with a cut-off score of 3.0-4.0. seqNLS (Lin et al., 2013) was used to determine which cilia proteins contain potential NLS sequences with a cut-off score of 0.3-0.5. The prediction scores across the three databases were normalized to give a z-score ranging from -2 to +2. Heatmap was generated using the heatmap.2 RStudio script. Candidate NLS-containing cilia proteins were selected based on NLS score and previously published roles in either ciliogenesis or cilia length regulation.

#### Microscopy

Images were acquired on an EVOS M7000 epifluorescent microscope or a Zeiss LSM 980 confocal microscope, as written in the figure legends. Images acquired on the EVOS M7000 were processed using the Celleste Imaging System 2D/3D Deconvolution at default settings. Images acquired on the Zeiss LSM 980 were acquired in either confocal mode or with Airyscan processing on Zen.

##### Cell culture

Importin α ciliary localization (Figure 1) and cilia length regulation (Figure 4) in RPE-1 cells were imaged with an Olympus Semi-Apo 60x 0.7 numerical aperture (NA) oil objective. U-ExM RPE-1 cell (Figure 1C) was imaged on a Zeiss 980 confocal microscope with a Plan Apo 63x, 1.4 NA oil objective. Drug treated and/or transfected RPE-1 cells (Figures 2-3) were imaged with an Olympus Semi-Apo 60x, 0.7 NA oil objective. Localization of importin α and Cep164 in RPE-1 cells (Figure 5) were imaged on EVOS M7000 epifluorescent microscope with an Olympus Super Apo 100x, 1.4 NA oil objective. Transfection of RPE-1 cells with Cep164 WT GFP or Cep164 ∆NLS GFP (Figure 5) were imaged on EVOS M7000 epifluorescent microscope with an Olympus Fluorite 20x, 0.45 NA air objective.

##### X. laevis

Importin α ciliary localization in epidermal MCCs (Figure 1) were imaged on EVOS M7000 epifluorescent microscope with an Olympus Super Apo 100x, 1.4 NA oil objective. Drug treated *X. laevis* NF stage 37 embryos (Figure 3B), *X. laevis* embryonic kidney images (Figures 6A-6B and Figure S5), and PORCN MO *X. laevis* embryos (Figures S2B and S2D) were imaged on EVOS M7000 epifluorescent microscope with an Olympus Semi-Apo 4x, 0.13 NA objective. Microinjected epidermal MCCS (Figure 3D) and PORCN MO epidermal MCCs (Figures S2C and S2E) were imaged on EVOS M7000 epifluorescent microscope with an Olympus Semi-Apo 60x, 0.7 NA oil objective. Reversible deciliation epidermal MCCs (Figure 4E) were imaged on EVOS M7000 epifluorescent microscope with an Olympus UPLFLN 40x, 0.51 NA air objective. Importin α and Cep164 localization at the epidermal MCC base was imaged on a Zeiss 980 confocal microscope with a Plan Apo 100x, 1.4 NA oil objective. X. laevis embryonic kidney, proximal and distal tubule were imaged on EVOS M7000 epifluorescent microscope with an Olympus Semi-Apo 60x, 0.7 NA oil objective.

### QUANTIFICATION AND STATISTICAL ANALYSIS

All statistical analyses were performed in GraphPad Prism 10.0. Comparisons between datasets were determined by a Student’s t-test unless otherwise stated. Graphs represent the mean value ± standard error of the mean (SEM) unless otherwise stated. *P<0.05, **P <0.01, ***P<0.001, and ****P<0.0001 unless otherwise stated. Blinding was not performed unless otherwise stated.

## Notes

### Competing Interest Statement

The authors have declared no competing interest.

