## Supplemental Figures for "A Role for Importin α in Ciliogenesis and Cilia Length Regulation during Nephrogenesis"

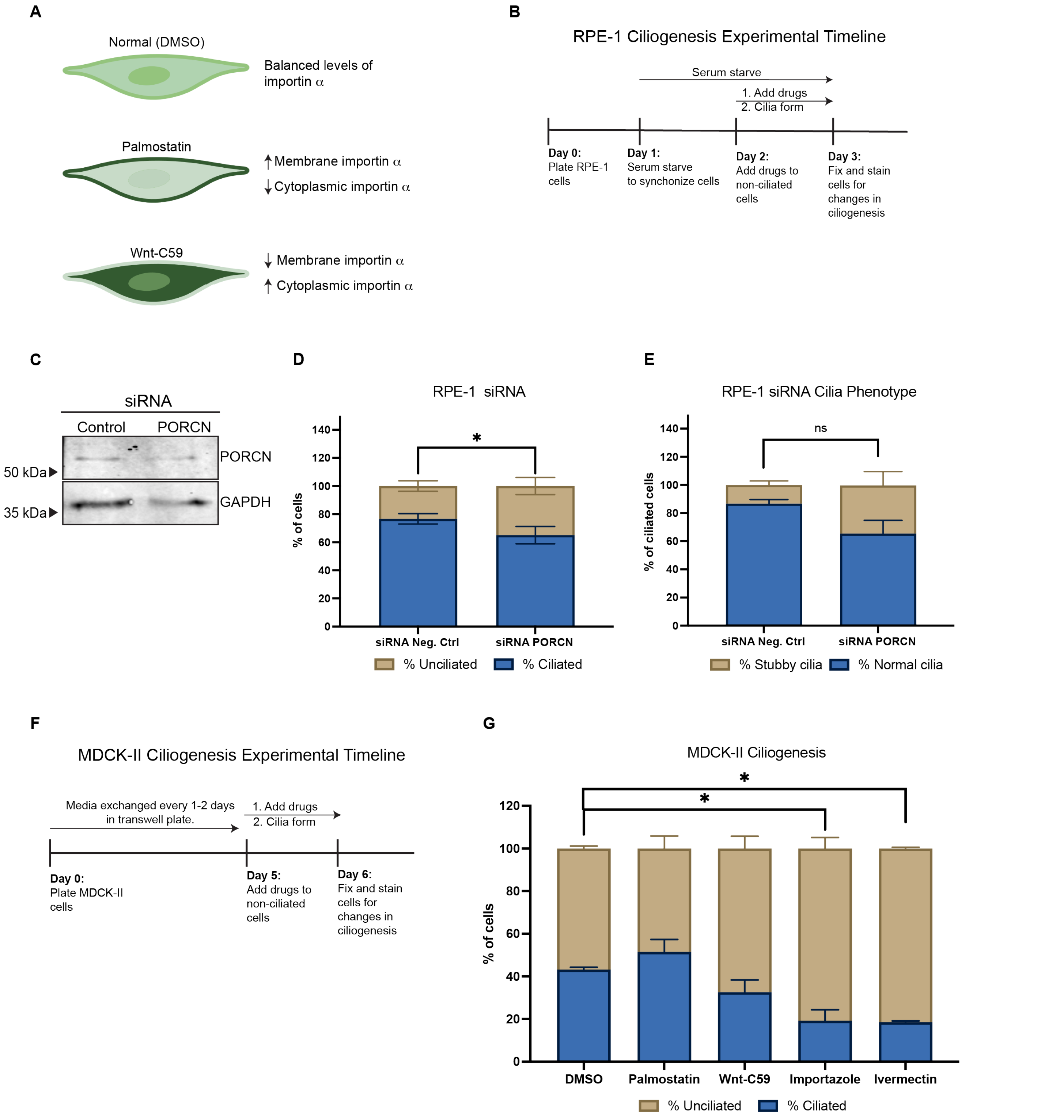


**Supplemental Figure 1: Palmitoylated importin α regulates the alternative ciliogenesis pathway. A)** Schematic of cellular and membrane levels of importin upon Wnt-C59 and Palmostatin treatment. **B)** Timeline for treating RPE-1 cells with drugs to inhibit palmitoylation status and importin function. **C)** Immunoblot of Porcupine O-acyltransferase (PORCN**)** knockdown via siRNA transfection in RPE-1 cells, immunoblotting for PORCN and GAPDH. **D)** Quantification of non-ciliated RPE-1 cells transfected with siRNA negative control (Neg. Ctrl) or siRNA PORCN for 24 hours and assessed for changes in the % of ciliated and % of unciliated cells. Student's t-test, * P<0.05, n= 250 cells per treatment, 3 replicates. Error bars are mean ± SEM. **E)** Quantification of % siRNA PORCN ciliated cells of having either a normal or stubby cilia phenotype. **F)** Timeline for treating non-ciliated MDCK-II cells with drugs to inhibit palmitoylation status and importin function. **G)** MDCK-II cells were treated with listed drugs for 24 hours: DMSO, 50µM palmostatin, 10µM Wnt-C59, 40µM importazole, or 25µM Ivermectin. Quantification of %ciliated vs. %unciliated MDCK-II cells after drug array treatment for 24hrs.* $P<0.05$, n= 750 cells. 2 replicates. Mean ± SEM.

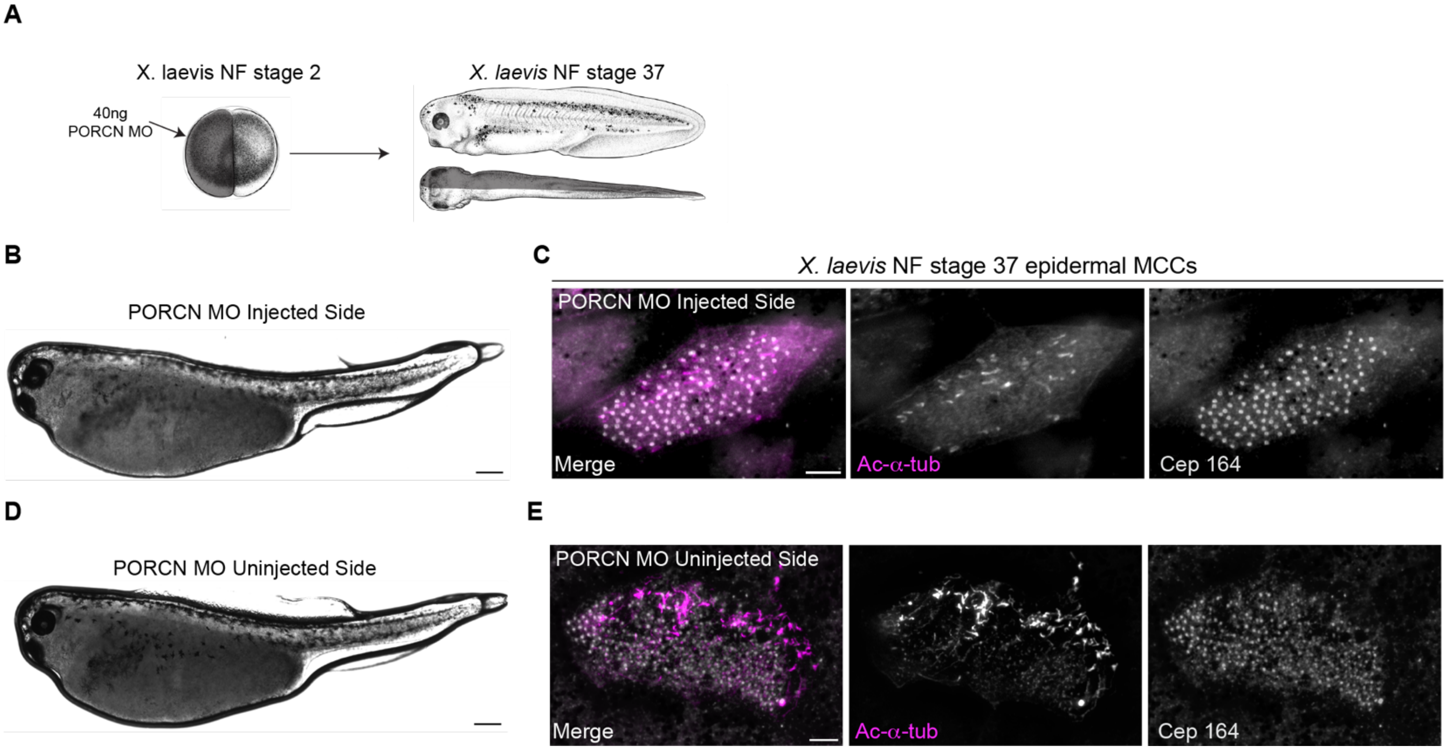


**Supplemental Figure 2: Disruption of multiciliogenesis in *X. laevis* epidermal MCCs using PORCN Morpholino (MO). A)** *X. laevis* NF stage 2 embryos were microinjected with 40ng of PORCN MO in one blastomere at the two-cell stage, targeting the epidermal MCCs. The uninjected blastomere serves as an internal control. **B)** Representative image of a PORCN MO injected half of a *X. laevis* NF stage 37 embryo. **C)** Epifluorescent image of an epidermal MCC from the PORCN MO-injected side immunostained for acetylated-α-tubulin (magenta) and Cep164 (gray). Scale bar = 200µm. **D)** Representative image of a PORCN MO uninjected half of a *X. laevis* NF stage 37 embryo. **E)** Epifluorescent image of an epidermal MCC from the uninjected side immunostained for acetylated-α-tubulin (magenta) and Cep164 (gray). Scale bar = 200µm.

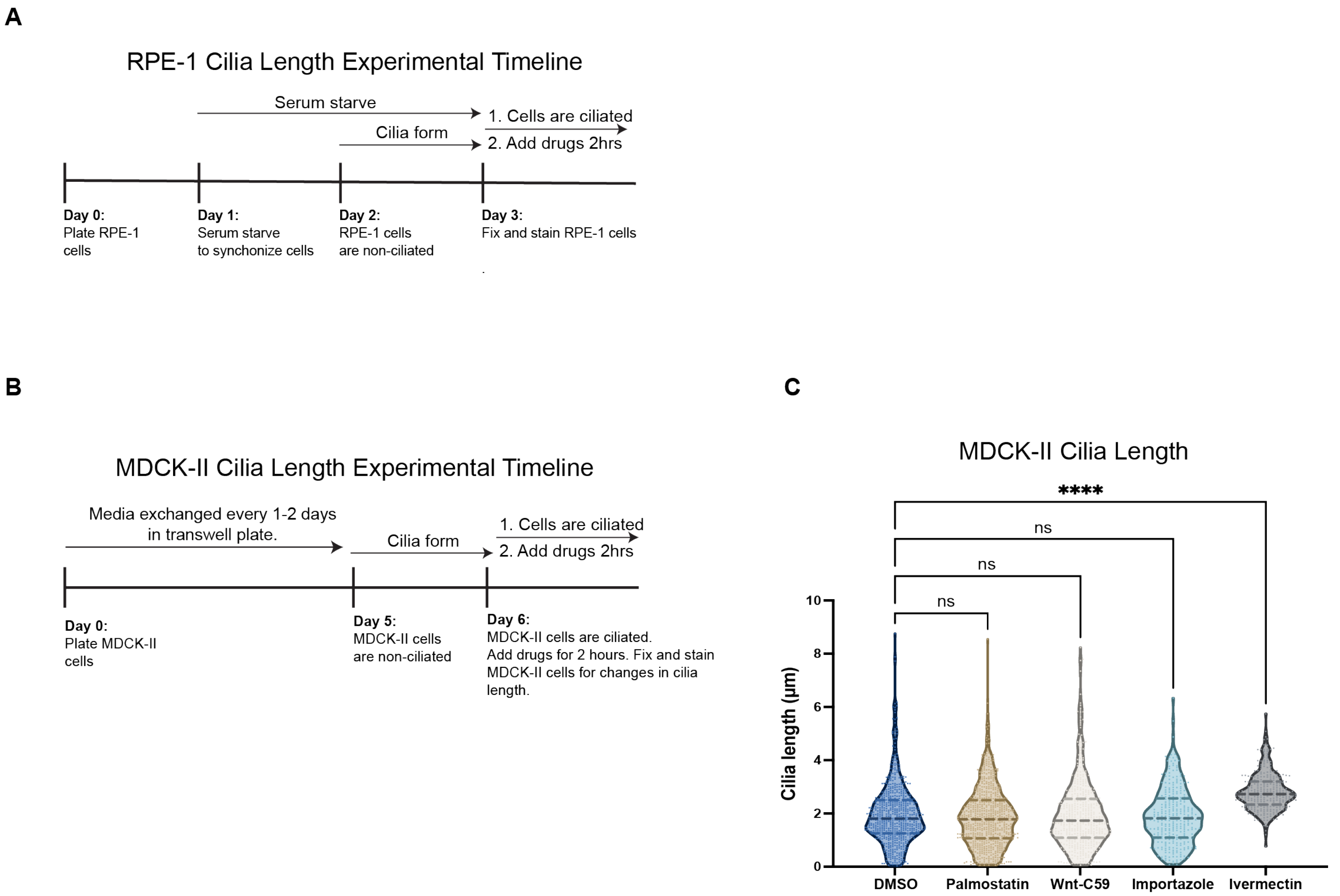


**Supplemental Figure 3: MDCK-II proper cilia length is regulated by palmitoylated importin α. A)** Timeline depicting serum starving RPE-1 cells for 48hrs in order to initiate ciliogenesis. Once ciliated, RPE-1 cells are treated with drugs to inhibit palmitoylation status and importin function for 2 hours with subsequent fixing and staining to quantify cilia length changes. **B)** Timeline of MDCK-II ciliated cells treated with drugs with subsequent monitoring of changes in cilia length. **C)** Quantification of cilia length (µm) of ciliated MDCK-II cells treated for 2hrs with drugs. ****$P<0.0001$ Student's t-test, n=750 cells per treatment, 2 replicates.

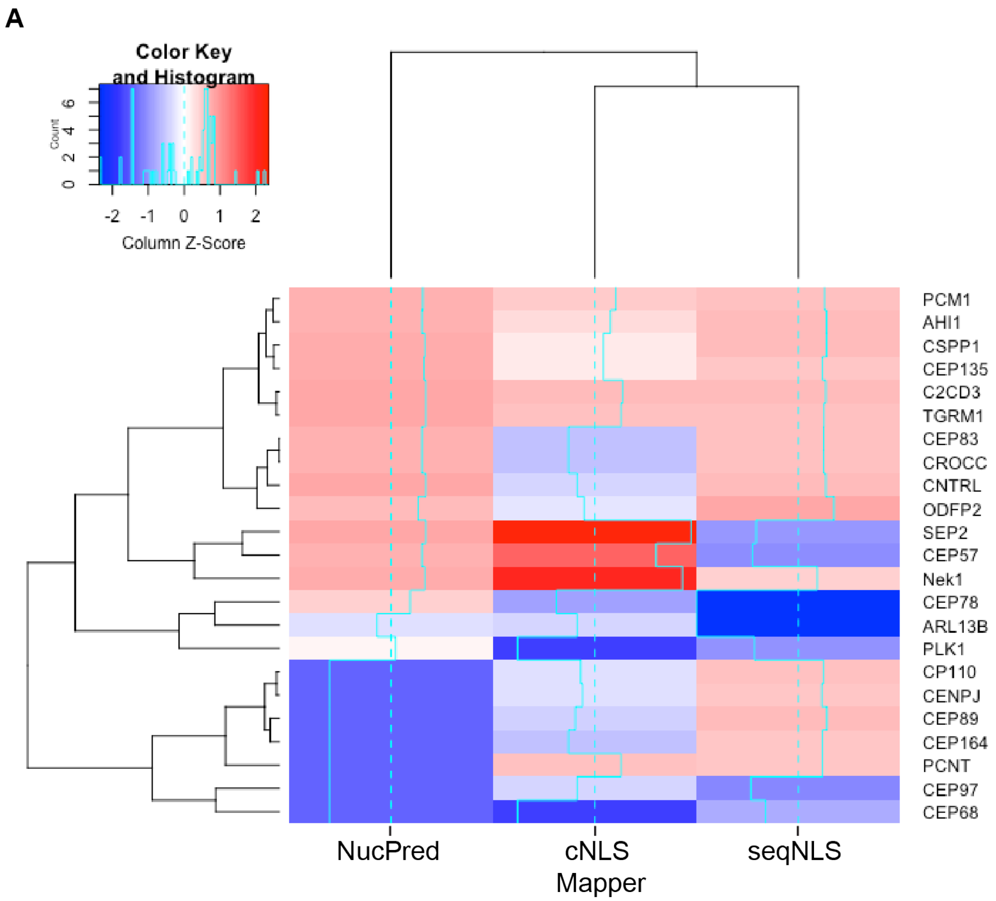


**Supplemental Figure 4: Ciliogenesis proteins that contain NLS sequences**. **A)** The amino acid sequence of proteins known to be involved in the early stages of ciliogenesis were systematically searched for a predicted NLS motif using three independently published NLS prediction tools; NucPred, cNLS Mapper, and seqNLS. NLS prediction scores from each database were normalized with a histogram with a ‘column Z-score’ (cyan). The ‘column Z-score’ indicates the NLS prediction score from -2 to 2. The Z-score for each cilia protein contains a trace from 0 (cyan dashed line) to show the NLS prediction score of containing a high probability of a true NLS sequence (red) or low probability of a true NLS sequence (blue).

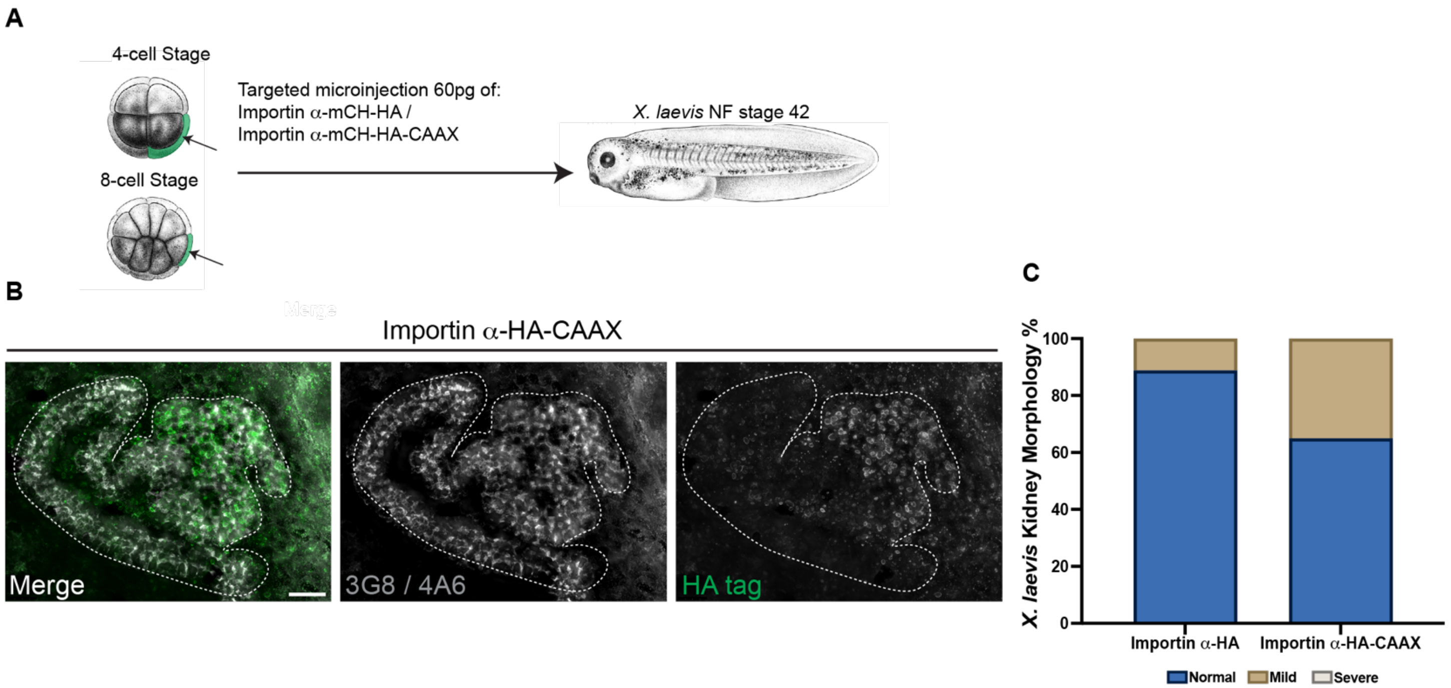


**Supplemental Figure 5: Forced membrane localization of importin α via CAAX motif does not impact kidney morphology. A)** *X. laevis* 4-cell or 8-cell embryos were microinjected with 60pg importin α-mCH-HA or importin α-mCH-HA-CAAX in blastomere highlighted in green to target the kidney. **B)** Deconvolved epifluorescent image of a microinjected NF stage 42 embryonic kidney with importin α-HA-CAAX plasmid, immunostained for kidney, 3G8/4A6 (gray) and HA-tag (green). **C)** Quantification of % *X. laevis* embryonic kidneys with normal, mild or severe kidney morphology after kidney-targeted microinjections of plasmids containing either importin α-mCH-HA or importin α-mCH-HA-CAAX. n≥9 tadpoles per microinjection.
